# Substrate binding and lipid-mediated allostery in the human organic anion transporter 1 at the atomic-scale

**DOI:** 10.1101/2022.07.14.500056

**Authors:** Angelika Janaszkiewicz, Ágota Tóth, Quentin Faucher, Hélène Arnion, Nicolas Védrenne, Chantal Barin-Le Guellec, Pierre Marquet, Florent Di Meo

**Affiliations:** Inserm U1248 Pharmacology & Transplantation, Univ. Limoges, 87000 Limoges, France; CHU de Tours, 2 Boulevard Tonnellé, 37044 Tours, France; Department of Pharmacology and Toxicology, CHU Limoges, F-87000 Limoges, France

**Keywords:** Membrane Transporters, Structural Pharmacology, Molecular Dynamics, Protein-lipid interactions, Major Facilitator Superfamily

## Abstract

The Organic Anion Transporter 1 is a membrane transporter known for its central role in drug elimination by the kidney. *h*OAT1 is an antiporter translocating substrate in exchange for *α*-ketoglutarate. The understanding of *h*OAT1 structure and function remains limited due to the absence of resolved structure of *h*OAT1. Benefiting from conserved structural and functional patterns shared with other Major Facilitator Superfamily transporters, the present study intended to investigate fragments of *h*OAT1 transport function and modulation of its activity in order to make a step forward the understanding of its transport cycle. *μs*-long molecular dynamics simulation of *h*OAT1 were carried out suggesting two plausible binding sites for a typical substrate, adefovir, in line with experimental observations. The well-known B-like motif binding site was observed in line with previous studies. However, we here propose a new inner binding cavity which is expected to be involved in substrate translocation event. Binding modes of *h*OAT1 co-substrate *α*-ketoglutarate were also investigated suggesting that it may binds to highly conserved intracellular motifs. We here hypothesize that *α*-ketoglutarate may disrupt the pseudo-symmetrical intracellular charge-relay system which in turn may participate to the destabilisation of OF conformation. Investigations regarding allosteric communications along *h*OAT1 also suggest that substrate binding event might modulate the dynamics of intracellular charge relay system, assisted by surrounding lipids as active partners. We here proposed a structural rationalisation of transport impairments observed for two single nucleotide polymorphisms, p.Arg50His and p.Arg454Gln suggesting that the present model may be used to transport dysfunctions arising from *h*OAT1 mutations.

**Highlights:**

- Adefovir has at least two binding pockets on *h*OAT1 in the outward-facing conformation.
- The highly conserved B-motif within MFS is strongly involved in substrate binding.
- *α*-Ketoglutarate binds to the intracellular domain of *h*OAT1 and destabilizes its OF conformation.
- The lipid membrane bilayer plays an active role in the allosteric communication between intracellular and extracellular domains of *h*OAT1.

**Graphical Abstract:** The present work (from left): (*i*) reveals binding modes of adefovir (top) and *α*-ketoglutarate (bottom) to *h*OAT1; (*ii*) maps Single Nucleotide Polymorphisms on outward-facing (top) and inward-facing (bottom) conformation of *h*OAT1; (*iii*) asses the allosteric effect of lipidic environment and presence of substrates.

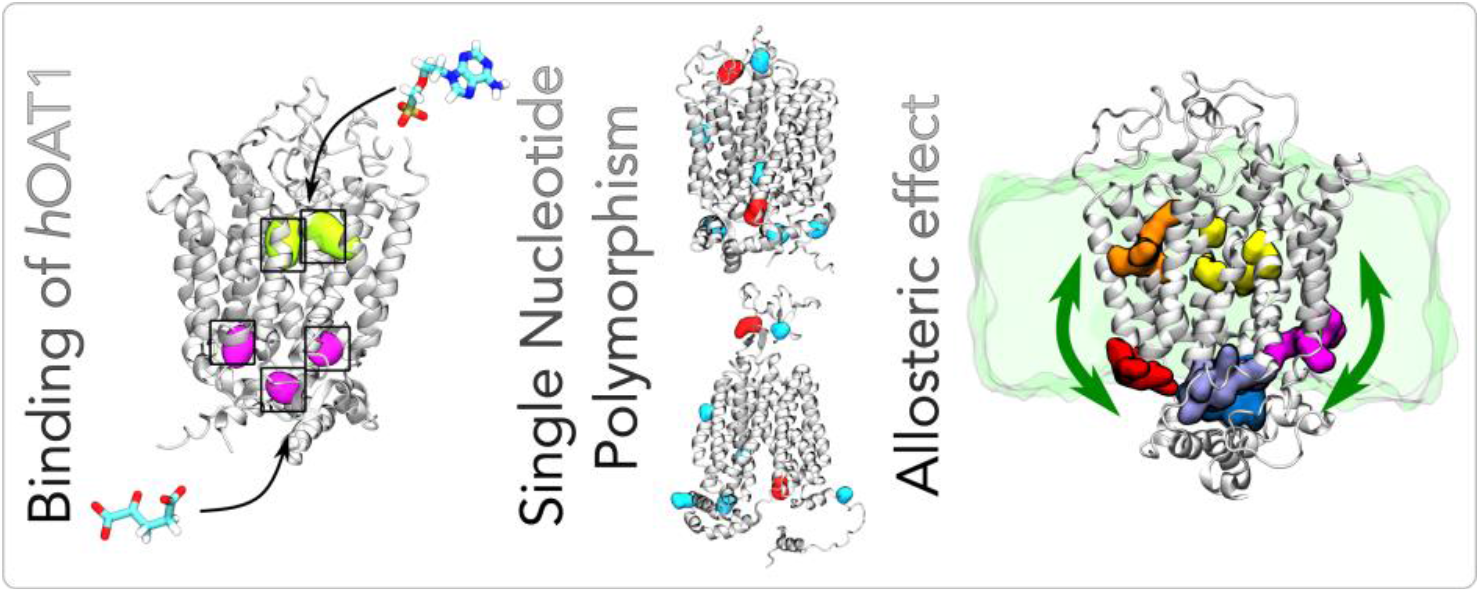

## 1. Introduction

Membrane transporters such as ATP-binding cassette (ABC) proteins and solute carriers (SLCs) are responsible for substrate translocation across cell membranes in a myriad of pharmacological and physiological events. At the cellular level, membrane transporters are involved in the selective influx and efflux of a broad range of compounds, including xenobiotics and endogenous substances. According to the “Remote Sensing and Signalling Theory” (RSST), which describes the transporter-mediated multi-organ regulatory system [1], the transepithelial transport of hormones [2], toxins [3], metabolites [4], and signalling molecules [1] at the interface between biofluids (e.g., blood/urine, blood/bile) plays an active role in body homeostasis. Pharmacological events such as drug absorption, distribution, metabolism, and elimination also involve membrane transporters. Local pharmacokinetics, i.e., drug concentration at the target sites, are directly governed by membrane transporters whether linked with drug therapeutic or adverse effects [5,6]. Understanding the role of membrane transporters in systemic drug pharmacokinetics is thus essential since defects in their expression and/or function might alter drug exposure, disposition, and response [1,6]. The International Transporter Consortium has listed transporters of “emerging clinical importance” for which the functional evaluations are recommended for new drug development [6,7]. The human organic anion transporter 1 (*SLC22A6*/*h*OAT1) belongs to this list given its central role in drug elimination as well as in the RSST [6–8]. As a key regulator of renal elimination, *h*OAT1 is expressed in the kidney proximal tubular cells (PTCs) at the basolateral membrane [9]. Diverse endogenous substrates, such as metabolites and signalling molecules, are transported by *h*OAT1. Additionally, xenobiotics like antiviral (such as adefovir), anticancer, and antituberculosis drugs are eliminated by *h*OAT1 [2].

*h*OAT1 is an antiporter, *i.e*., the uptake of substrates from the blood into PTCs is coupled to the excretion of *α*-ketoglutarate (*α*KG) as a co-substrate into blood circulation [10]. This requires a high intracellular concentration of *α*KG in PTCs. That is maintained thanks to the so-called tertiary active transport model in which *h*OAT1 activity is coupled with Na^+^/K^+^-ATPase pump and NaDC3/*SLC13A3* transporter [2,11]. Atomic-scaled mechanism of *h*OAT1-mediated transport remains unclear owing to the lack of experimentally resolved structure. Structural patterns of *h*OAT1 might provide insights into its roles regarding drug efficacy, toxicity, and elimination [12,13]. We recently proposed a structural and dynamic model of *h*OAT1 embedded in four different lipid bilayers models made of 1-palmitoyl-2-oleoyl-*sn*-glycero-3-phosphatidylcholine (POPC), 1-palmitoyl-2-oleoyl-*sn*-glycero-3-phosphatidylethanolamine (POPE) and cholesterol (Chol), namely POPC, POPC:Chol (3:1), POPC:POPE (3:1) and POPC:POPE:Chol (2:1:1) [14]. *h*OAT1 is expected to adopt the Major Facilitator Superfamily (MFS) fold for which substrate translocation follows the alternating access model. It requires large-scale conformational changes between at least two major conformational states, namely outward-facing (OF) or inward-facing (IF) conformations responsible for substrate binding and release events, respectively [15]. *h*OAT1 structure encompasses 12 transmembrane helices (TMH) organized into N- (TMH1-6) and C-bundles (TMH7-12). The transmembrane domain is formed by the functional A-, B-, and C-helices that appear as pseudo-symmetrical repetitions in *h*OAT1 structure [14,15]. Briefly, the inner cavity is made of A-helices (TMH1, 4, 7 and 10), which are expected to bind substrates and structurally adjust along the transport cycle. At the bundle interface, B-helices (TMH2, 5, 8, 11) support the large conformational changes along transport cycle. Finally, the out-of-the-core C-helices (TMH3, 6, 9 and 12) were suggested to maintain the structural integrity of the transporter through interactions with surrounding lipids [16]. Between TMH1 and TMH2, *h*OAT1 exhibits a large extracellular loop (ECL) with four identified glycosylation sites. ECL was suggested to be involved in trafficking; however, its glycosylation is non-essential for the transport function [17,18]. The intracellular region is involved in regulations of transport function and expression on the opposite side of the membrane [18,19]. The intracellular region was suggested to consist of 6 intracellular helices that are tightly connected to TMH intracellular regions. The presence of charged amino acids in conserved motifs is suggested to represent an MFS proteins signature. The so-called A-, E[X_6_]R and PETL motifs are pseudo-symmetrically repeated in both N- and C-bundles. The interactions between these motifs differ between IF and OF conformations [14]. We recently proposed that membrane bilayer components may interact with *h*OAT1 in specific exhibit lipid binding sites driven by H-bond and electrostatic interactions [14]. Phosphatidylethanolamine (PE), which was suggested to stabilise the OF conformation by its interaction with B-helices on the extracellular site, drew particular attention. This was in line with joint experimental and computational investigations suggesting lipid components might facilitate the transport cycle in MFS proteins [20].

Knowledge about binding modes of *h*OAT1 substrates remains fragmented, despite their importance for the understanding and prediction of transporter-mediated drug-drug interactions (DDIs) [21–23]. Likewise, investigating the structural impact of single nucleotide polymorphisms (SNPs) in *h*OAT1 may help to understand the mild transport impairments of *h*OAT1 observed experimentally, even though their clinical impacts may be rather limited owing to substrate overlap with other PTC influx transporters (e.g., *h*OAT3). The present study aimed to model the binding modes of adefovir, an acyclic nucleoside phosphonate (ANP) antiviral, to *h*OAT1. Adefovir was chosen as a *h*OAT1 substrate prototype given the extensive experimental data available in the literature [2,24,25]. To better understand the interplay with co-substrate translocation, attention was paid to the binding of co-substrate *α*KG on the intracellular region as well. We here also investigate the possible active role of lipid components in the distant communication between predicted adefovir binding sites and the charge-relay system since it had previously been mentioned as being essential for other MFS proteins [20,26,27].

## 2. Methods

### 2.1. Binding of (co-)substrates to *h*OAT1 model

From our previous μs-scaled MD simulations performed on the apo *h*OAT1 OF model, representative snapshots were extracted from MFS core based Principal Component Analysis (Figure S1) to investigate the binding modes of *α*KG and adefovir as co-substrate and substrate, respectively. Initial poses for (co-)substrates were obtained by means of molecular docking calculations using AutoDock Vina 1.1.2 [28]. *α*KG and adefovir were modelled as dianionic compounds given the physiological pH. *α*KG and adefovir initial structures were obtained from PubChem database [29] and optimized by quantum mechanical methods at the (CPCM)-M06-2X/6-31+G(d,p) level of theory using the Gaussian16 Rev. A package [30]. The partial charges and final parameters were obtained using antechamber [31,32] (the parameter files available in supplemental materials). Molecular docking search volumes were defined with 66560 and 64768 Å^3^ for the *h*OAT1 extracellular and intracellular sides respectively for adefovir and *α*KG given the antiporter transport (see Table S1 and Figure S2 for the definitions of box size and centre). For each substrate, 20 molecular docking calculations were carried out providing 20 molecular poses each, leading to 400 poses per substrate for each membrane. For each substrate, three initial binding poses were selected accounting for the calculated affinity scores as well as for the experimentally-identified key binding residues (see Figure S3) [33–35]. To confirm their relevance, molecular docking poses were then used as initial positions for μs-scaled MD simulations.

Three different systems were considered for further MD simulations, namely *α*KG-, adefovir- and *α*KG-adefovir-bound *h*OAT1. For each system, three binding modes were considered for further MD simulations (see Figure S3). In total, nine *h*OAT1 systems were used for MD simulations (see Figure S3). It is worth mentioning that two *α*KG molecules were systematically considered for *α*KG-based systems in order to optimize sampling. All systems were embedded into three types (POPC)-based membranes namely POPC, POPC:Chol (3:1) and POPC:POPE:Chol (2:1:1). In total, 27 different systems were considered in the present study. Systems were all solvated with water using a 0.154M NaCl concentration to mimic extracellular physiological conditions. System compositions and sizes are reported in supporting information (Table S2). It is important to note that one position (namely, position 3 in Figure S3) was excluded since adefovir left the binding cavity in all simulations, except in POPC:POPE:Chol (2:1:1) membrane. However, in these simulations, adefovir mostly interacted with lipid components at the lipid-protein-water interface rather than with *h*OAT1 residues.

### 2.2. MD simulation setup

For the definitions of the protein, lipids and water, the following forcefields were applied Amber FF14SB [36], Lipids17 [37] and TIP3P [38], respectively. The parameters for counterions (Na^+^, Cl^-^) were obtained from Joung and Cheatham [39,40]. The definition of the co-substrate and substrate required a parametrization using GAFF2 and DNA.OL15 forcefield parameters, considering the latter being a purine derivative. Charges were derived using the Antechamber package available in the Amber18 package [41,42], considering QM-optimized structures at the density functional theory level (namely M062X/6-31+G(d,p)) in implicit water using the conductor-like polarizable continuum model. MD simulations were performed using a similar setup as previously established [14]. The simulations were carried out with using the CPU and GPU codes of the Amber18 package [41,42]. The system was treated with periodic boundary conditions. Non-covalent interactions were considered within the cut-off of 10 Å for electrostatic and Lennard-Jones potentials. Long-range electrostatic interactions were treated using the particle mesh Ewald (PME) method [43]. The bonds involving hydrogen atoms were fixed by applying the SHAKE algorithm. Integration time step was set at 2 fs. To maintain the physiological conditions, the temperature was set at 310K and carried on using a Langevin thermostat [44]. Whereas the Monte Carlo barostat was applied to maintain the constant pressure boundary condition under semi-isotropic conditions [45].

The equilibration of the system was pursued by minimizing all atomic positions and followed by a smooth two-step thermalization. The system was heated up from 0 to 100 K during 200 ps under (N, V, T) conditions, while the second step of the thermalization up to 310K was carried out under semi-isotropic (N, P, T) conditions. Subsequently, equilibration simulations were performed for 5.5 ns. Finally, the MD simulations of substrate-bound *h*OAT1 resulted in a duration of 1.5 μs each, leading to a total of *ca*. 40.5 μs. Trajectory snapshots were saved every 10 ps.

### 2.3. Analysis

Given the previous validation of the model [14], structural analyses were based on our recent findings, as well as on the general knowledge of MFS patterns [15]. Structural analyses were performed using the PyTRAJ and CPPTRAJ AMBER modules [46] as well as the VMD software [47]. Based on the time-dependent backbone root-mean squared deviations (Figure S4), analyses were performed on the equilibrated section of the present trajectories, i.e., over the last 800 ns (Figure S4). H-bond analyses were performed using distance and angle cutoffs set at 3.0 Å and 135°, respectively. H-bond analyses were rationalized considering the fraction of frames the H-bond is present, accounting the number of H-bond. For instance, two residues interacting along in every single frame thanks to 1 or 2 H-bond(s) will lead to H-bond fraction values at 1.0 and 2.0, respectively. The minimum fraction threshold to discard irrelevant H-bonds was set at 0.1 given the known uncertainties for side chain rotameric states in protein threading techniques. MD trajectories obtained from our previous study was also used for sake of comparison between apo and bound *h*OAT1 systems.

To unravel the allosteric communications between key regions of *h*OAT1, we carried out network analyses using Allopath tool [48]. The communication efficiency between two distant domains was calculated from the protein contact map and the mutual information matrix in which the contributions of interactors (*e.g*., substrates and lipids) were included. Communication efficiencies were calculated unidirectionally from “source” to “sink” residues (see Ref. [48] for more details). In the present work, allosteric communications between binding pocket residues (B-like motif and the inner binding cavity, see Section 3.3) and the charge-relay system (A- and E[X_6_]R motifs, see Table S3 and Section 3.3) were calculated. The contributions of interactors were assessed by calculating communication efficiencies on systems made of: (i) *h*OAT1 *apo*, (ii) *h*OAT1 substrate-bound, (iiI) *h*OAT1 *apo* with lipids, and (iv) *h*OAT1 substrate-bound with lipids.

## 3. Results and Discussion

### 3.1. Binding sites and key residues of *h*OAT1 revealed through interactions with (co-)substrates

#### 3.1.1. Adefovir binds to B-motif and inner binding cavity of *h*OAT1

*h*OAT1 was shown to play a key role in the uptake of anionic endogenous and exogenous substrates, such as antiviral acyclic nucleoside phosphonates (ANP) (*e.g*., tenofovir, adefovir) [35,49], urate, *p*-aminohippurate (PAH), β-lactam antibiotics and sulfate conjugates [50].

The presence of numerous cationic residues (*i.e*., lysine, arginine, and histidine, see Figure 1A) in intracellular and extracellular interfaces of IF and OF *h*OAT1 models is consistent with the expected high affinity for anionic substrates by favouring electrostatic interactions. Cationic residues were also found in the ECL between TMH1 and TMH2 as well as in the water-exposed cavity. Molecular docking calculations initially suggested three possible binding site for adefovir as well as for *α*KG in the intracellular regions (Figure S3). The adefovir binding site (position 3 in Figure S3) was however excluded from MD simulations since adefovir spontaneously left the predicted binding site. MD simulations confirmed the importance of A-helices in the water-exposed cavity, mainly TMH1 and TMH4. The latter was previously reported as key for substrate binding and translocation owing to the presence of the conserved so-called B-like motif RXX[Q/S]G [51]. The B-like motif was observed in several MFS antiporters, even though it cannot be considered as an antiporter fingerprint [51]. MD simulations reveal that adefovir may strongly bind to the B-like motif Arg192 residue thanks to a salt-bridge between phosphonate and guanidinium moieties as pictured by calculated large H-bond fraction at 1.950, where the maximum value can be 2.0 over the MD simulation (see Figure 1B,C and Table S4). H-bond analyses also highlighted the key role of several polar and aromatic residues, namely Ser255, Gln251, Ser195 and Tyr141. The stability of the substrate was enhanced by *π*-stacking interactions with Tyr141 between the adefovir adenosyl moiety and tyrosine phenol moiety. This is in line with the *h*OAT1 substrate spectrum which includes various anionic aromatic compounds (*e.g*., ANPs, PAH, urate). Interestingly, the *h*OAT1 OF model indicates that there are two possible access pathways to the B-motif cavity. Substrates may enter directly from the water phase through the ECL cationic residue network or an access channel between TMH3 and TMH6 that may exist, connecting the high-density polar head region of the outer leaflet membrane with the B-like motif cavity. This is consistent with biophysical studies showing that amphiphilic substrates might partition beneath polar head region of the lipid bilayer membrane [52].

**Figure 1.**
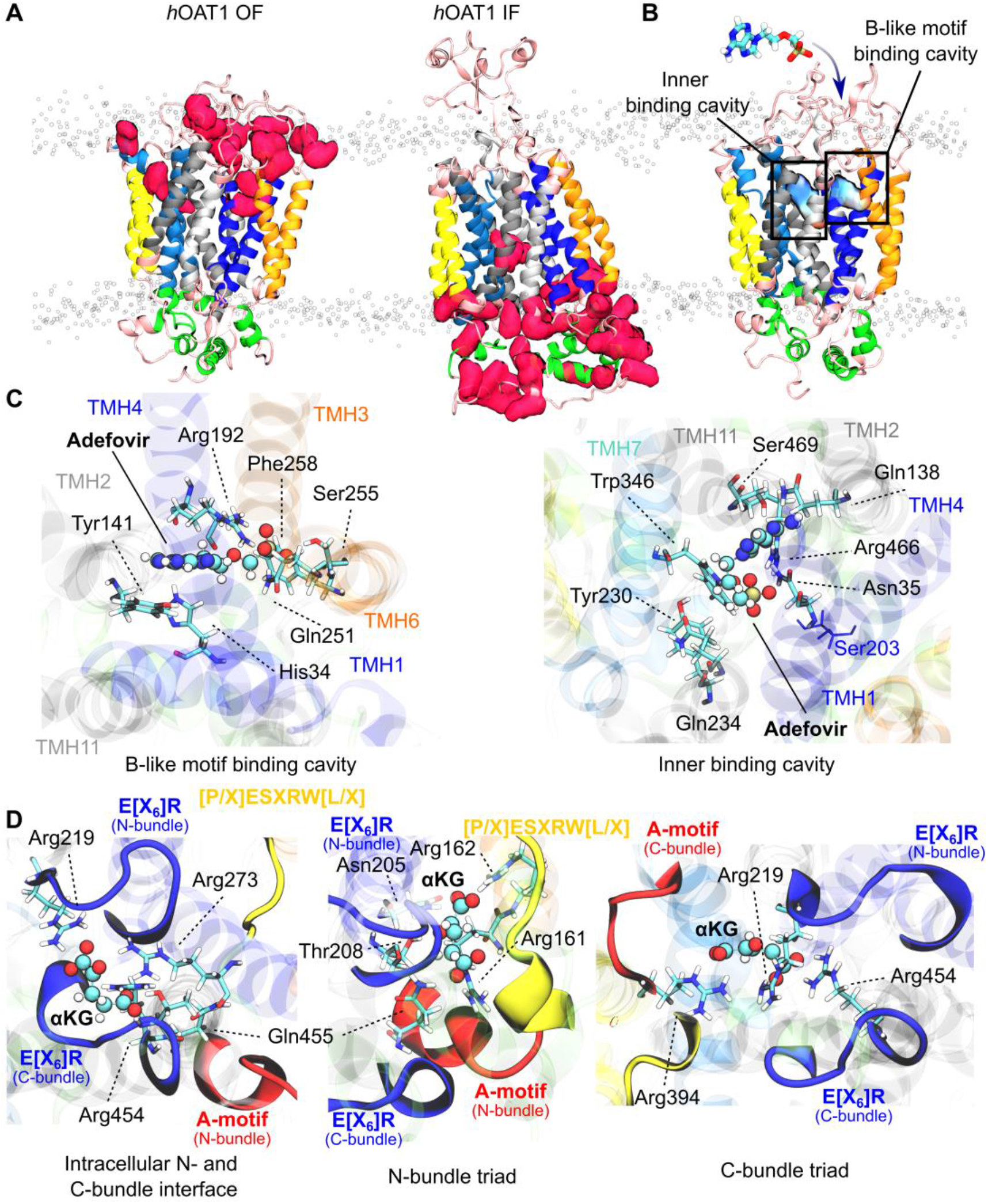
Interactions with (co-)substrate. A) Representation of cationic residues (pink spheres) present in extracellular and intracellular sites of the OF (left) and IF (right) models embedded in lipid bilayer (PC and PE P- and N-atoms being depicted in white spheres). B) Adefovir is absorbed by the water-exposed cavity of *h*OAT1 OF model, which results in 2 binding pockets: the B-like motif and the inner binding cavity involving TMH1 and TMH4 (dark blue). C) Residues involved in adefovir binding in the B-like motif (left) and inner (right) binding cavity. D) Three mains *α*KG binding spots and the most frequently interacting residues. For all panels, model is coloured according to A- (light and dark blue), B- (light and dark grey) and C- (yellow and orange) transmembrane helices colour code.

The existence of an inner binding cavity was observed at the interface between N- and C-bundles, involving mostly TMH1 and TMH7. The substrate is again stabilized thanks to a strong H-bond network (Figure 1D). Cationic Arg466 is likely to play a central role for anionic substrate translocation (Table S4 and Figure 1C). This is in good agreement with experimental observations in which site-directed mutagenesis of Arg466 was associated to substrate-dependent loss of transport capacity (Table S5) [53]. The first- or second-shell residues around adefovir contain several of site-directed mutagenesis residues that were associated to the loss of transport function (*e.g*., Asn39 [17], Tyr230 [33], Tyr353 [54], Trp346 [54], see Table S5). This confirms the relevance of the region obtained from MD-refined molecular docking calculations. Particular attention was also paid to Ser203 (Figure 1C) which was described as central for the binding of several ANPs. In our simulations, Ser203 belongs to the second-shell of contact residues with adefovir. Consequently, no direct interactions between adefovir and Ser203 were observed in our calculations. Binding to Ser203, however, was suggested from static structural investigations performed on an IF *h*OAT1 homology model [35]. It may indicate that Ser203 is likely engaged in substrate translocation by transporting substrate from the outer to the inner leaflet as part of the OF-to-IF large-scale conformational transition, which is consistent with experimental observations. This is further strengthened by the recent AlphaFold2 IF model [55] in which Ser203 is deeply located in the MFS core, precluding direct substrate binding in the OF conformation.

The inner binding cavity is slightly less accessible than the B-like motif pocket, likely due to the absence of an access channel. Interestingly, the two cavities are contiguous, around TMH1. Therefore, present results as well as experimental observations regarding mutations of residues located in the inner binding cavity [33,34,53,54] suggest that substrate binding might occur first in the B-like motif cavity. Substrate binding to B-like motif pocket is thus expected to rapidly lead to local conformational changes which may open an access channel to the inner binding cavity. Binding to inner cavity may is in turn be pivotal for substrate translocation events along the transport cycle. Finally, rocker-switch large-scale TMH conformational changes occurring during the translocation event might decrease substrate binding affinity, favouring substrate release in the intracellular medium.

#### 3.1.2. α-Ketoglutarate may disrupt the conserved network of intracellular interactions destabilizing the OF conformation

*h*OAT1 transport cycle requires large conformational changes for the OF-to-IF transition. *h*OAT1 being an antiporter, substrate uptake is coupled with the co-transport of *α*KG in the opposite direction [10]. However, the sequence of events (*i.e*., *α*KG efflux and substrate influx) remains unclear as well as triggers. Even though our present model cannot provide a clear solution to that issue, it can be used to propose binding modes of *α*KG in the intracellular region of *h*OAT1. As stated for adefovir binding at the extracellular interface, intracellular domains are also rich in cationic residues favouring electrostatic interactions with anionic *α*KG molecules. These cationic residues are exposed to the intracellular medium independently on the conformational state (Figure 1A and 1B). Even though *α*KG translocation is expected to reset the OF conformation, our simulations suggest that *α*KG can also bind intracellular domains, regardless of the *h*OAT1 conformational state.

Three preferential binding modes were obtained for *α*KG: (i) on the N- and (ii) C-bundle motif triads; or (iii) at the interface between N- and C-bundle E[X_6_]R motifs (Figure 1D). Owing to its two anionic carboxylate moieties, *α*KG largely favours electrostatic interactions with charge-relay system cationic residues, as pictured by described H-bond network from MD simulations (Figure 1D, Tables S4 and S6). For instance, strong salt-bridges were observed along the simulations between *α*KG and A-motif arginine residues, namely Arg161 and Arg162, or Arg394 for binding poses in the N- or C-bundle triad, respectively (H-bond fractions being 2.019, 1.698 and 1.376 for Arg162, Arg161 and Arg394, respectively). Likewise, the intracellular charge-relay system might play a central role in *α*KG binding at the interface between N- and C-bundle E[X_6_]R motifs (e.g., Arg219, Arg454). In turn, MD simulations with *α*KG bound system to the intracellular domain also reveal that the charge-relay system might be disrupted by the presence of *α*KG (Table S6). Assuming that *α*KG can bind *h*OAT1 in the OF conformation, present results suggest that *α*KG may play favour substrate translocation by destabilizing intracellular N- and C-bundle interactions, which is necessary for OF-to-IF transition along with substrate translocation. These findings pave the way for further investigations to establish whether the IF-to-OF transition resetting *h*OAT1 conformation is driven by *α*KG efflux or if *α*KG and the other substrate are simultaneously transported; even though the latter is expected to be less likely [15].

### 3.2. The lipid environment is largely responsible for the increase in allosteric communication between the substrate binding pockets and intracellular domains

The comparison of the intracellular structural arrangement between OF and IF apo *h*OAT1 models revealed that the charge-relay system must be disrupted along transport cycle. This might ultimately unlock the IC gating required for substrate release [14]. Conformational changes are expected to be triggered upon substrate and/or co-substrate binding, suggesting the existence of a distant communication between substrate binding pockets and MFS conserved intracellular motifs. Allosteric communications were thus monitored providing structural insights about the efficiency regarding the different domains located across the membrane (see Section 2.3 and Ref. [48] for technical details). Particular attention was paid to the plausible communications between binding domains (B-like motif and inner substrate binding site) with the charge-relay system (A- and E[X_6_]R motifs). It is worth mentioning that (i) N- and C-bundle IC motifs were considered separately and (ii) allostery was monitored in both directions, *i.e*., from EC to IC regions and *vice versa* leading to 16 different pathways (Figures 2 and S5-S11). The role of substrate, co-substrate and surrounding lipid bilayer were also considered. However, results provided here are only qualitatively discussed since the resolution of the present model, especially regarding side chains, precludes quantitative conclusions.

**Figure 2.**
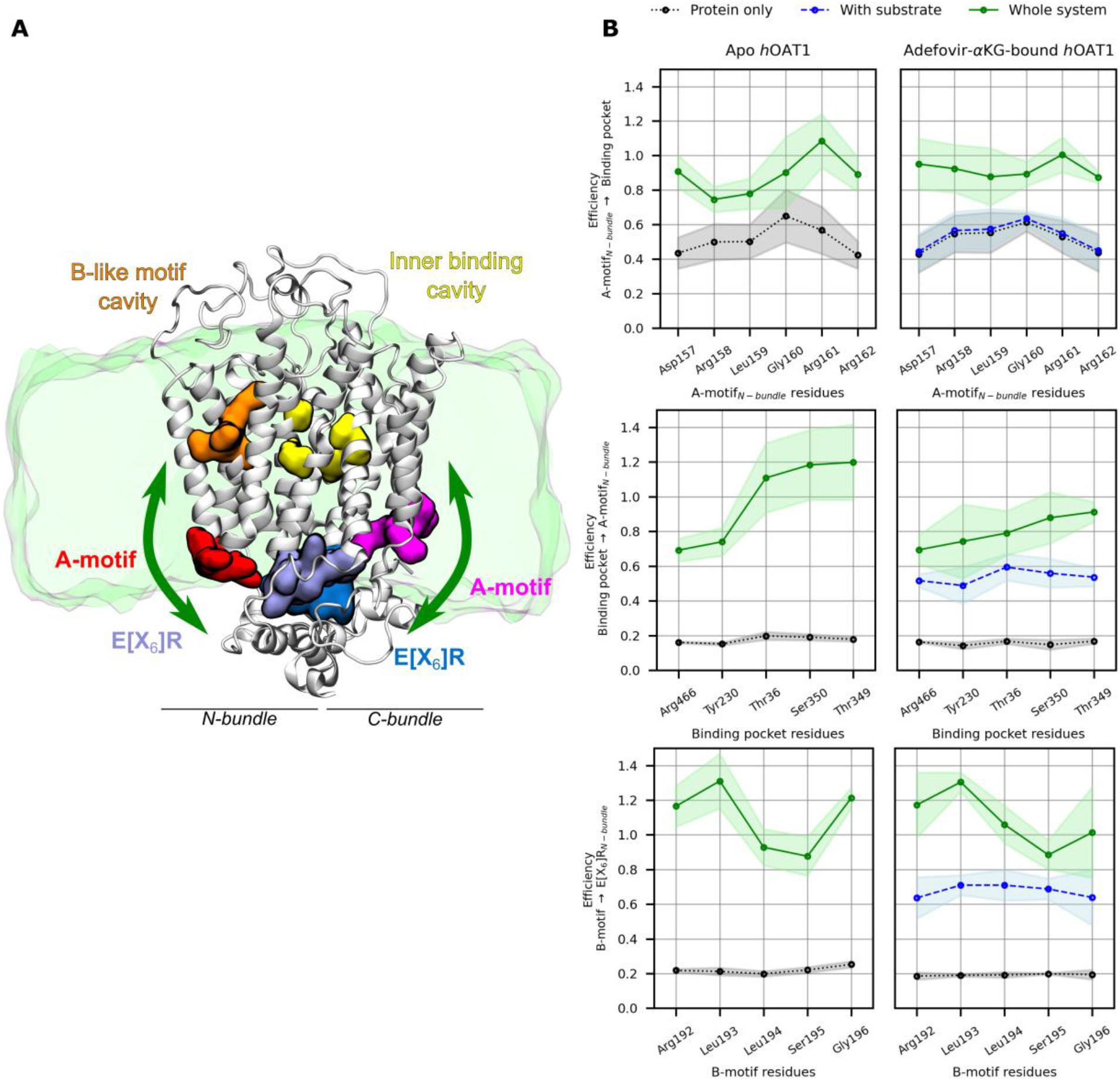
The allosteric effect of the membrane lipid bilayer in the presence and absence of substrates on the *h*OAT1 communication efficiency from biding pockets toward charge-relay system and reverse. (A) Visualisation of binding pockets (B-like motif cavity on left and inner binding cavity on right) and motifs of charge-relay system divided into N and C domain. The wide green arrows represent strong communication in presence of lipids. (B) The schemes plot the communication efficiency calculated for pure protein (black dashed line), protein in presence of substrates (blue dashed line) and protein accounting lipids in apo (left column) or substrate-bound state (right column) in green line.

As an example, Figure 2B shows efficiencies of selected allosteric pathways, namely between N-bundle A-motif and inner binding pocket in the two directions as well as between B-like motif and N-bundle E[X_6_]R motif. The existence of a reciprocal distant communication between intracellular motifs and substrate binding pocket was observed from MD simulations. While considering only the role of *h*OAT1 protein, allosteric pathway analyses revealed an efficient communication from A-motifs to substrate binding pockets, while other pathways remain inherently low. Furthermore, these allosteric pathway efficiencies exhibit very similar profiles for both B-like motif and inner binding pockets. Interestingly, efficiencies calculated for allosteric pathways from A-motifs to substrate binding pockets were not affected by the presence of substrates. In contrast, the allosteric communications from binding pockets to intracellular motifs significantly potentiated the efficiency in the presence of substrate and/or co-substrate. In other words, substrate binding events are thus expected to modulate the dynamics of intracellular motifs. Such observations are in line with the proposed mechanism of *h*OAT1 transport for which binding events might trigger the conformational changes in the intracellular region required for substrate translocation. The involvement of *h*OAT1 residues (or betweenness) to allosteric networks were also calculated as shown in Figure S9-S12. These analyses underlined the pivotal roles of TMH2, TMH3 and TMH4 which may be easily explained since they are part of binding pockets as well as in the N-bundle A-motif.

Besides, it is important to note that MFS protein functions have been experimentally shown to strongly depend on the membrane composition [20,26,56]. Likewise, PE lipids were shown to non-covalently bind with the residues standing at the interface between N- and C bundles in apo *h*OAT1 structure [14]. As a consequence of the structural interplay between surrounding lipid bilayer and *h*OAT1 protein, the active role of lipid bilayer membrane in the communication between extracellular and intracellular regions was also observed. Efficiencies drastically increased while considering the membrane. This confirms that the membrane plays an active role in allosteric signalling as recently shown for other membrane proteins [48], including transporters [57]. This also underlines the central role of protein-lipid interactions and the need for representative and realistic membranes, including PE and cholesterol, for computational and experimental investigations on *h*OAT1. At the body level, we can thus expect that *in situ* alteration or modulation of cell membrane composition may affect the activity of *h*OAT1 either by modifying the global structure and dynamics of *h*OAT1, or its intrinsic function [58,59]. Such observations may be extended to other MFS transporters given the high protein-lipid dependency.

### 3.3. Structural mapping of *h*OAT1 single nucleotide polymorphisms contributes to understanding the transport impairment caused by meaningful mutations

Beyond pharmacologically relevant mechanistic insights into *h*OAT1-mediated drug transport, the present structural model can also be used to understand the modulation of substrate transport across cell membranes by naturally-occurring polymorphisms in genes coding transporters [60–63]. Several SNPs in *h*OAT1 were identified and classified according to ethnical origins and species [35]. Since most SNPs are located in untranslated intronic regions, the focus will be given here to non-silent mutations (Table 1 and Figure 3), which correspond to a limited number of SNPs reported in the literature. Moreover, most of them exhibit no or limited alteration of transport function. The present IF and OF *h*OAT1 models can help understand the structural changes induced by genetic polymorphisms.

**Table 1.**
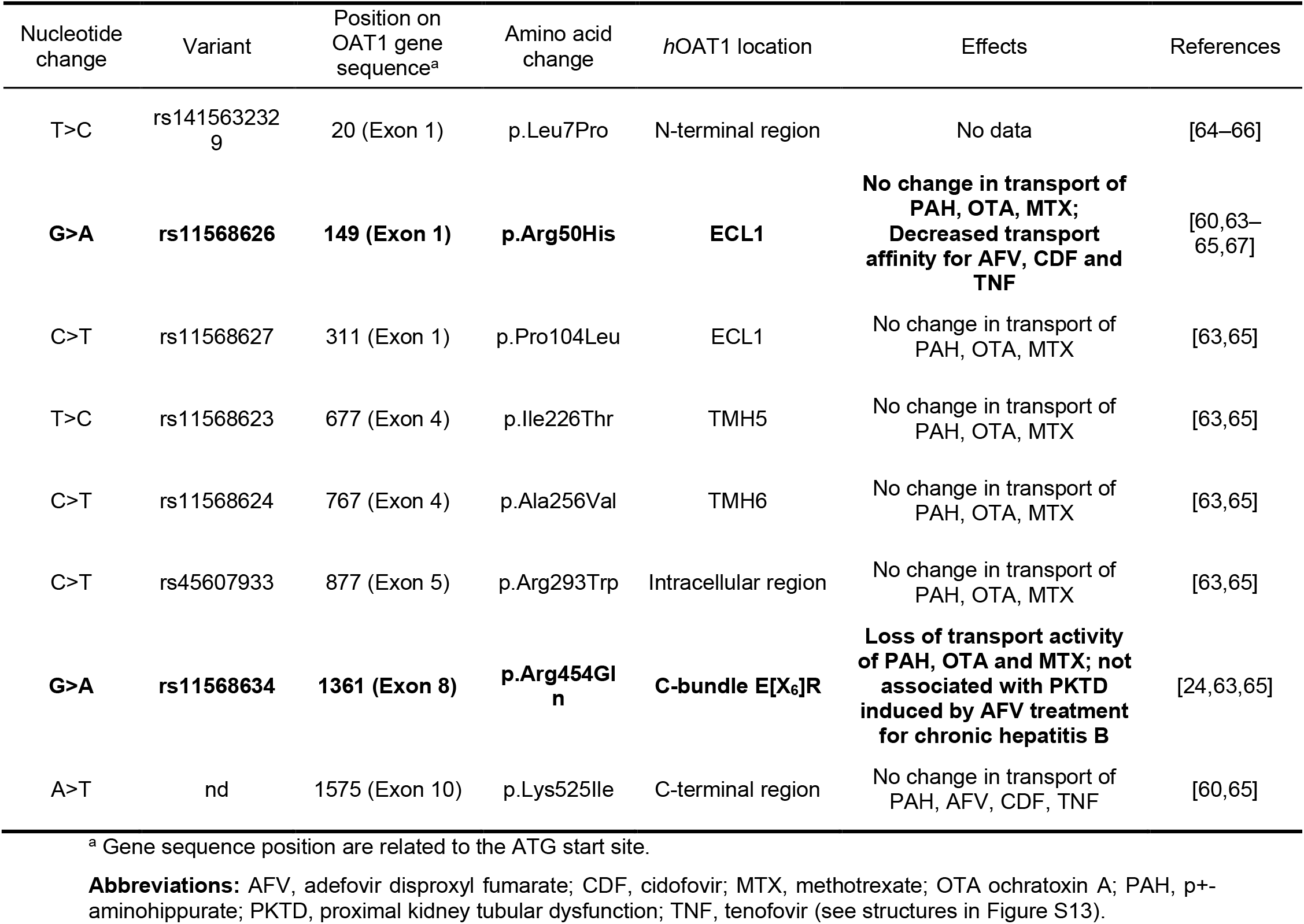
List of *h*OAT1 SNPs and rare variants. The reported impairment of *h*OAT1 function is shown in bold.

**Figure 3.**
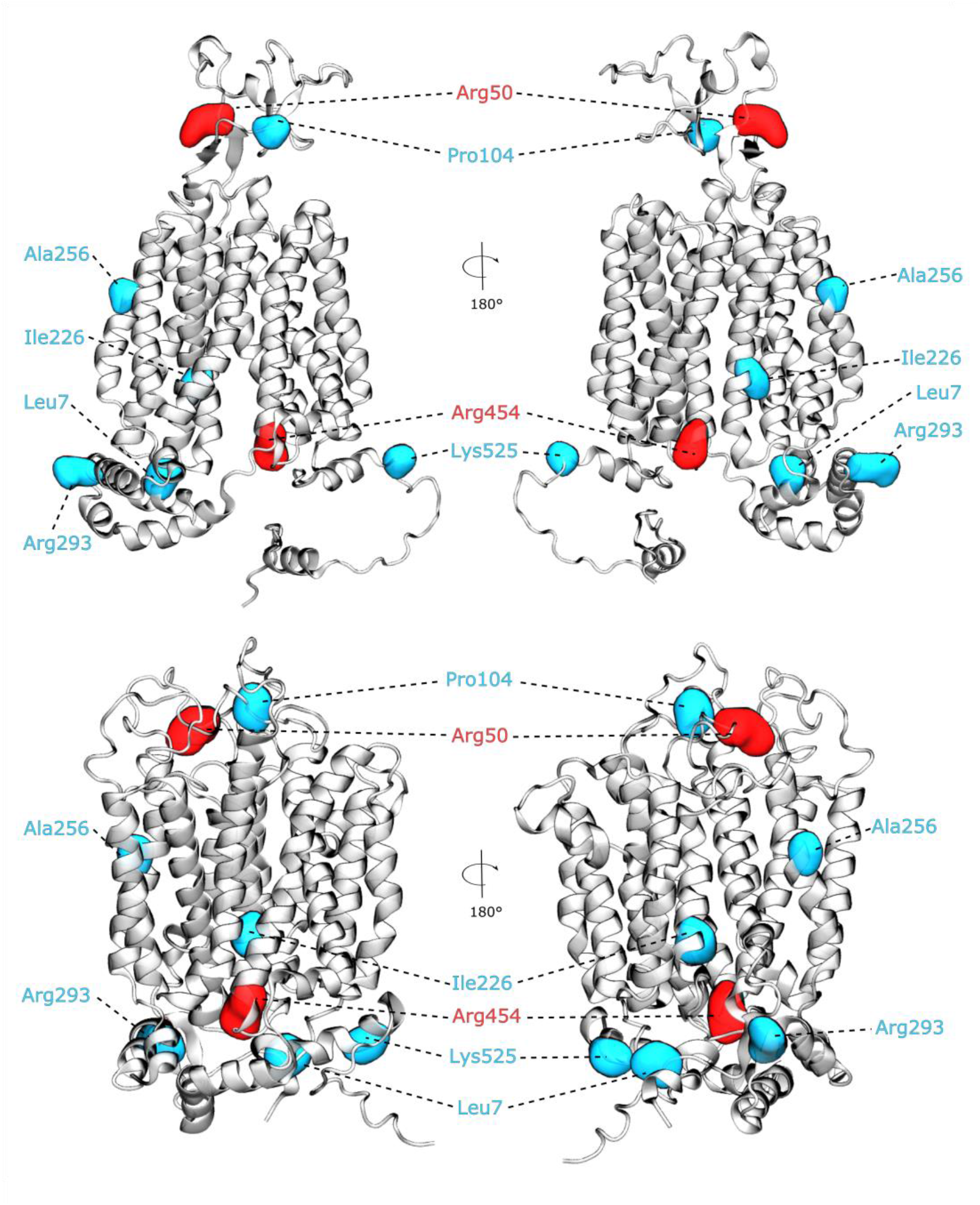
Mapped Single Nucleotide Polymorphisms on IF (top) and OF (bottom) conformations of *h*OAT1. The position of mutated residues associated with a partial loss of functions are depicted in red (Arg50 and Arg454) while those for which no impact was reported are coloured in cyan (Pro104, Ala256, Arg293, Ile226 and Leu7).

Particular attention will be paid to those that were experimentally suggested to affect transport activity rather than protein expression or altered membrane trafficking. Eight SNPs have been considered and mapped on the present IF and OF *h*OAT1 models (Figure 3). They can be classified according to their location on the model, as follows: (i) in the N-terminal domain (p.Leu7Pro); (ii) in the long ECL between TMH1 and TMH2 (p.Arg50His and p.Pro104Leu); (iii) in TMH5 and TMH6 (p.Ile226Thr and p.Ala256Val); and (iv) in the intracellular domain (p.Arg293Trp, p.Arg454Gln and p.Leu525Ile). SNPs leading to p.Pro104Leu, p.Ile226Thr, p.Ala256Val, p.Arg293Trp and p.Lys525Ile substitutions were not experimentally associated with alteration of substrate intake [24,60,63,64,67]. The proposed structural models of hOAT1 in IF and OF conformations can thus be used to propose explanation regarding the absence of change in transport function. Ala256 and Ile226 are located in transmembrane domain, at the interface of lipid bilayer membrane, and are not expected to be involved in key regions involved in (co -)substrate translocation event nor in folding issues. This is particularly true the rs11568624 variant for which Ala256 is mutated into valine, maintaining hydrophobic nature of the residue. Pro104 is located in the flexible ECL between TMH1 and TMH2 which has been mostly suggested to be involved in post-translational modifications (glycosylation) [10, 17]. We can thus hypothesise that proline mutation may not significantly affect the dynamics nor the electrostatic potential of ECL1, the latter being more important for anionic substrate binding event. On the other side of the membrane, Arg293 is located in the intracellular domain, but our models suggest that it is spatially distant from charge-relay system. Therefore, its mutation into tryptophane (rs45607933) is thought to not strongly affect the intracellular domain dynamics.

Only two SNPs were reported to partially impair *h*OAT1 function, namely rs11568626 and rs11568634 leading to p.Arg50His and p.Arg454Gln protein mutations, respectively. The rs11568626 SNP was found to be specific to the African population. The resulting p.Arg50His mutation was shown to be associated with decreased transport affinities (K_m_) of phosphate analogues such as adefovir, cidofovir and tenofovir in Xenopus oocytes-expressed p.Arg50His variant as compare to wild-type *h*OAT1 (rs15914676). However, Xenopus oocytes-expressing p.Arg50His variant exhibited normal uptake of PAH, ochratoxin A and methotrexate [63]. Interestingly, Arg50 is located in a conserved motif of the long ECL between TMH1 and TMH2, rich in cationic residues and the present model suggests that ECL is likely involved in anionic substrate access to the B-like motif binding pocket. We can hypothesize that the p.Arg50His substitution may lead to a lower electrostatic potential between the anionic substrate and the binding access channel.

This would also smooth the observations suggesting that the ECL suggesting a may not be involved in transport function of hOAT1 [17,18]. Given the poor ECL resolution of the present OF model, this must be confirmed by further experiments or through the experimental structure resolution of *h*OAT1. Interestingly, another SNP involving a residue located in ECL was reported (rs11568627, p.Pro104Leu) but no transport impairment was observed. This might be by explained by the neutral feature of such mutation, *i.e*., neutral proline to neutral leucine, which does not modulate the aforementioned electrostatic potential. The rs11568634 SNP leads to the substitution of Arg454 by a glutamine residue located on the other side of the membrane,. Interestingly, this SNP was associated to decreased uptake of PAH, ochratoxin A and methotrexate, suggesting a loss of function [63]. This is in perfect agreement with present structural observations which underlined the central role of Arg454 in the intracellular charge-relay system: Arg454 is likely involved in salt-bridges between the C-bundle A-motif and the PETL motif. The p.Arg454Gln substitution is thus expect to weaken the local supramolecular arrangement, which in turn is likely to disrupt the charge-relay system dynamics essential for the MFS transport cycle.

It is worth mentioning that no *h*OAT1 SNP was associated with pathological conditions at the clinical level. Owing to the central role of OATs in RSST [1], the substrate overlap between *e.g*., *h*OAT1 and *h*OAT3 [10] might lead to compensation activity between transporters. However, systemic compensation between transporters might not be sufficient to overcome *h*OAT1 impairment in chronic conditions such as chronic kidney diseases. Recently, particular attention has been paid to the loss of tubular function in chronic kidney disease [68], where *h*OAT1 impairment is expected to play a key role [69]. Further clinical investigations are required to examine *h*OAT1 SNPs as risk factors for the still unclear patient variability in *e.g*., long-term ANP nephrotoxicity [25,60,63,70].

## 4. Conclusion

The present study investigated substrate binding events of both substrate and co-substrate to the human *SLC22A6*/OAT1 at the atomic scale. Molecular docking calculations and μs-scaled MD simulations revealed two plausible binding spots for adefovir, consistent with experimental observations, in the B-like motif and inner binding pockets. A-helices (essentially TMH1, TMH4) residues were shown to play an essential role for substrate binding event. In the intracellular region, MD simulations also suggested the binding of *α*KG involved residues of the charge-relay system, located either within the motif triads (within A-motif, E[X_6_]R and PETL) or at the interface between them. The presence of *α*KG may interfere with the salt-bridge network of intracellular conserved motifs. This is expected to favour the opening of IC gating which might be key in driving large-scale conformational changes required for *h*OAT1 alternated access. The allosteric analysis was performed to demonstrate the communication between the substrate binding sites and intracellular motifs, which supported the hypothesis that substrate binding may indirectly affect large conformational changes by opening the intracellular gate. When only the protein was taken into account, TMH2, 3, and 4 played a major role in the allosteric signal’s pathway. However, as lipids and (co-)substrates are also considered in allosteric communication, the involvement of TMH2, 3, and 4 was drastically reduced. Lipids consistently increased allosteric communication, regardless of the presence of (co-)substrates. This demonstrates the significance of lipids as a component of the allosteric signal pathway and, consequently, their crucial role for the transport function of *h*OAT1. Non-synonymous SNPs were then mapped onto the *h*OAT1 structural model. Transport impairments experimentally observed for p.Arg50His and p.Arg454Gln were respectively attributed to (i) the decrease of substrate binding affinity or (ii) disrupted intracellular domain interactions owing to the weakening of electrostatic interactions. Each new SNP or site-direct mutagenesis mutant can now be implemented in the present model to elucidate its role at an atomistic resolution.

Altogether, the present study paves the way for the structural understanding of *h*OAT1 function in a pharmacological context. The present model can be used to understand and predict transporter-xenobiotic interactions as well as DDIs. The allosteric pathways described may be involved in non-competitive DDIs, by which a *h*OAT1 inhibitor may interact with an allosteric site involved in the communication between key regions.

## Supporting information

Electronic Supplementary information

## Abbreviations

ABC: ATP-Binding Cassette
AFV: Adefovir
*a*KG: *α*-ketoglutarate
ANP: Acyclic Nucleoside Phosphonate
CDF: Cidofovir
Chol: Cholesterol
DDI: Drug-drug interaction
ECL: ExtraCellular Loop
H-bond: Hydrogen bond
IC: IntraCellular
IF: Inward-Facing
K_M_: Binding affinity
MD: Molecular Dynamics
MFS: Major Facilitator Superfamily
MTX: Methotrexate
NaDC3: Na^+^/dicarboxylate transporter
OAT: Organic Anion Transporter
OF: Outward-Facing
OTA: Ochratoxin A
PAH: *p*-aminohippurate
PE: PhosphatidylEthanolamine
PKTD: Proximal Kidney Tubular Dysfunction
PME: Particle Mesh Ewald
POPC: 1-palmitoyl-2-oleoyl-sn-glycero-3-phosphocholine
POPE: 1-palmitoyl-2-oleoyl-sn-glycero-3-phosphoethanolamine
PTC: Proximal Tubular Cells
RSST: Remote Sensing Signalling Theory
SLC: SoLute Carrier
SNP: Single Nucleotide Polymorphisms
TMH: TransMembrane Helix
TNF: Tenofovir

## 6. CRediT (Contributor Role Taxonomy) author contributions

**Angelika Janaszkiewicz:** Conceptualization, methodology, formal analysis, investigation, writing (original draft, reviewing & editing). **Ágota Tóth:** Methodology, validation, formal analysis, investigation, writing (reviewing & editing). **Quentin Faucher:** Conceptualization, formal analysis, investigation, writing (reviewing & editing). **Hélène Arnion:** formal analysis. **Nicolas Védrenne:** formal analysis, writing (reviewing & editing). **Chantal Barin-Le Guellec:** Conceptualization, supervision, writing (reviewing & editing). **Pierre Marquet:** Conceptualization, supervision, funding acquisition, writing (reviewing & editing). **Florent Di Meo:** Conceptualization, methodology, validation, formal analysis, investigation, writing (original draft, reviewing & editing), supervision, funding acquisition.

## 7. Acknowledgments

This work was granted access to the HPC resources of IDRIS under the allocations 2020 - A0080711487, 2021-A0100711487, 2022-A0120711487 made by GENCI, using the GPU supercomputer “Jean Zay”. We are grateful to regional supercomputers CALI (“CAlcul en LImousin”) & “Baba Yaga”. Authors are grateful to Xavier Montagutelli for technical support as well as to Marving Martin and Dr. Benjamin Chantemargue and Dr. Marie Essig for fruitful scientific discussions.

## 9. Supplementary Information

In Supplementary information are reported: (i) molecular docking and MD technical details; (ii) calculated H-bond and contact fractions; (iii) list of site-directed mutagenesis reviewed from the literature; (iv) Allosteric communication efficiencies for all systems. Data are available upon reasonable request, including MD trajectories, Force field parameters and inputs/scripts used for MD simulations and allopath tool.

## References

[1] S.K. Nigam, What do drug transporters really do?, Nat. Rev. Drug Discov. 14 (2015) 29–44. https://doi.org/10.1038/nrd4461.

[2] A.L. VanWert, M.R. Gionfriddo, D.H. Sweet, Organic anion transporters: discovery, pharmacology, regulation and roles in pathophysiology, Biopharm. Drug Dispos. (2009) n/a-n/a. https://doi.org/10.1002/bdd.693.

[3] D.H. Sweet, Organic anion transporter (Slc22a) family members as mediators of toxicity, Toxicol. Appl. Pharmacol. 204 (2005) 198–215. https://doi.org/10.1016/j.taap.2004.10.016.

[4] Q. Faucher, H. Alarcan, P. Marquet, C. Barin-Le Guellec, Effects of Ischemia-Reperfusion on Tubular Cell Membrane Transporters and Consequences in Kidney Transplantation, J. Clin. Med. 9 (2020) 2610. https://doi.org/10.3390/jcm9082610.

[5] R. Ho, R. Kim, Transporters and drug therapy: Implications for drug disposition and disease, Clin. Pharmacol. Ther. 78 (2005) 260–277. https://doi.org/10.1016/j.clpt.2005.05.011.

[6] The International Transporter Consortium, Membrane transporters in drug development, Nat. Rev. Drug Discov. 9 (2010) 215–236. https://doi.org/10.1038/nrd3028.

[7] M.J. Zamek-Gliszczynski, M.E. Taub, P.P. Chothe, X. Chu, K.M. Giacomini, R.B. Kim, A.S. Ray, S.L. Stocker, J.D. Unadkat, M.B. Wittwer, C. Xia, S.-W. Yee, L. Zhang, Y. Zhang, International Transporter Consortium, Transporters in Drug Development: 2018 ITC Recommendations for Transporters of Emerging Clinical Importance, Clin. Pharmacol. Ther. 104 (2018) 890–899. https://doi.org/10.1002/cpt.1112.

[8] K.M. Hillgren, D. Keppler, A.A. Zur, K.M. Giacomini, B. Stieger, C.E. Cass, L. Zhang, Emerging Transporters of Clinical Importance: An Update From the International Transporter Consortium, Clin. Pharmacol. Ther. 94 (2013) 52–63. https://doi.org/10.1038/clpt.2013.74.

[9] H.C. Liu, N. Jamshidi, Y. Chen, S.A. Eraly, S.Y. Cho, V. Bhatnagar, W. Wu, K.T. Bush, R. Abagyan, B.O. Palsson, S.K. Nigam, An Organic Anion Transporter 1 (OAT1)-centered Metabolic Network, J. Biol. Chem. 291 (2016) 19474–19486. https://doi.org/10.1074/jbc.M116.745216.

[10] S.K. Nigam, K.T. Bush, G. Martovetsky, S.-Y. Ahn, H.C. Liu, E. Richard, V. Bhatnagar, W. Wu, The Organic Anion Transporter (OAT) Family: A Systems Biology Perspective, Physiol. Rev. 95 (2015) 83–123. https://doi.org/10.1152/physrev.00025.2013.

[11] M. Roth, A. Obaidat, B. Hagenbuch, OATPs, OATs and OCTs: the organic anion and cation transporters of the SLCO and SLC22A gene superfamilies: OATPs, OATs and OCTs, Br. J. Pharmacol. 165 (2012) 1260–1287. https://doi.org/10.1111/j.1476-5381.2011.01724.x.

[12] L. Wang, D.H. Sweet, Renal Organic Anion Transporters (SLC22 Family): Expression, Regulation, Roles in Toxicity, and Impact on Injury and Disease, AAPS J. 15 (2013) 53–69. https://doi.org/10.1208/s12248-012-9413-y.

[13] S.K. Nigam, W. Wu, K.T. Bush, M.P. Hoenig, R.C. Blantz, V. Bhatnagar, Handling of Drugs, Metabolites, and Uremic Toxins by Kidney Proximal Tubule Drug Transporters, Clin. J. Am. Soc. Nephrol. 10 (2015) 2039–2049. https://doi.org/10.2215/CJN.02440314.

[14] A. Janaszkiewicz, Á. Tóth, Q. Faucher, M. Martin, B. Chantemargue, C. Barin-Le Guellec, P. Marquet, F. Di Meo, Insights into the structure and function of the human organic anion transporter 1 in lipid bilayer membranes, Sci. Rep. 12 (2022) 7057. https://doi.org/10.1038/s41598-022-10755-2.

[15] D. Drew, R.A. North, K. Nagarathinam, M. Tanabe, Structures and General Transport Mechanisms by the Major Facilitator Superfamily (MFS), Chem. Rev. 121 (2021) 5289–5335. https://doi.org/10.1021/acs.chemrev.0c00983.

[16] E.M. Quistgaard, C. Löw, F. Guettou, P. Nordlund, Understanding transport by the major facilitator superfamily (MFS): structures pave the way, Nat. Rev. Mol. Cell Biol. 17 (2016) 123–132. https://doi.org/10.1038/nrm.2015.25.

[17] K. Tanaka, W. Xu, F. Zhou, G. You, Role of glycosylation in the organic anion transporter OAT1, J. Biol. Chem. 279 (2004) 14961–14966. https://doi.org/10.1074/jbc.M400197200.

[18] J. Zhang, H. Wang, Y. Fan, Z. Yu, G. You, Regulation of organic anion transporters: Role in physiology, pathophysiology, and drug elimination, Pharmacol. Ther. 217 (2021) 107647. https://doi.org/10.1016/j.pharmthera.2020.107647.

[19] C. Zhu, K.B. Nigam, R.C. Date, K.T. Bush, S.A. Springer, M.H. Saier, W. Wu, S.K. Nigam, Evolutionary Analysis and Classification of OATs, OCTs, OCTNs, and Other SLC22 Transporters: Structure-Function Implications and Analysis of Sequence Motifs, PLOS ONE. 10 (2015) e0140569. https://doi.org/10.1371/journal.pone.0140569.

[20] C. Martens, M. Shekhar, A.J. Borysik, A.M. Lau, E. Reading, E. Tajkhorshid, P.J. Booth, A. Politis, Direct protein-lipid interactions shape the conformational landscape of secondary transporters, Nat. Commun. 9 (2018) 4151. https://doi.org/10.1038/s41467-018-06704-1.

[21] A. Gessner, J. König, M.F. Fromm, Clinical Aspects of Transporter-Mediated Drug–Drug Interactions, Clin. Pharmacol. Ther. 105 (2019) 1386–1394. https://doi.org/10.1002/cpt.1360.

[22] Y. Liang, S. Li, L. Chen, The physiological role of drug transporters, Protein Cell. 6 (2015) 334–350. https://doi.org/10.1007/s13238-015-0148-2.

[23] X. Huo, K. Liu, Renal organic anion transporters in drug–drug interactions and diseases, Eur. J. Pharm. Sci. 112 (2018) 8–19. https://doi.org/10.1016/j.ejps.2017.11.001.

[24] M. Shimizu, N. Furusyo, H. Ikezaki, E. Ogawa, T. Hayashi, T. Ihara, Y. Harada, K. Toyoda, M. Murata, J. Hayashi, Predictors of kidney tubular dysfunction induced by adefovir treatment for chronic hepatitis B, World J. Gastroenterol. 21 (2015) 2116–2123. https://doi.org/10.3748/wjg.v21.i7.2116.

[25] C.C. Wong, N.P. Botting, C. Orfila, N. Al-Maharik, G. Williamson, Flavonoid conjugates interact with organic anion transporters (OATs) and attenuate cytotoxicity of adefovir mediated by organic anion transporter 1 (OAT1/SLC22A6), Biochem. Pharmacol. 81 (2011) 942–949. https://doi.org/10.1016/j.bcp.2011.01.004.

[26] C. Martens, R.A. Stein, M. Masureel, A. Roth, S. Mishra, R. Dawaliby, A. Konijnenberg, F. Sobott, C. Govaerts, H.S. Mchaourab, Lipids modulate the conformational dynamics of a secondary multidrug transporter, Nat. Struct. Mol. Biol. 23 (2016) 744–751. https://doi.org/10.1038/nsmb.3262.

[27] M.P. Muller, T. Jiang, C. Sun, M. Lihan, S. Pant, P. Mahinthichaichan, A. Trifan, E. Tajkhorshid, Characterization of Lipid–Protein Interactions and Lipid-Mediated Modulation of Membrane Protein Function through Molecular Simulation, Chem. Rev. 119 (2019) 6086–6161. https://doi.org/10.1021/acs.chemrev.8b00608.

[28] O. Trott, A.J. Olson, AutoDock Vina: Improving the speed and accuracy of docking with a new scoring function, efficient optimization, and multithreading, J. Comput. Chem. (2009) NA-NA. https://doi.org/10.1002/jcc.21334.

[29] PUBCHEM, (n.d.). https://pubchem.ncbi.nlm.nih.gov.

[30] M.J. Frisch, G.W. Trucks, H.B. Schlegel, G.E. Scuseria, M.A. Robb, J.R. Cheeseman, G. Scalmani, V. Barone, G.A. Petersson, H. Nakatsuji, X. Li, M. Caricato, A.V. Marenich, J. Bloino, B.G. Janesko, R. Gomperts, B. Mennucci, H.P. Hratchian, J.V. Ortiz, A.F. Izmaylov, J.L. Sonnenberg Williams, F. Ding, F. Lipparini, F. Egidi, J. Goings, B. Peng, A. Petrone, T. Henderson, D. Ranasinghe, V.G. Zakrzewski, J. Gao, N. Rega, G. Zheng, W. Liang, M. Hada, M. Ehara, K. Toyota, R. Fukuda, J. Hasegawa, M. Ishida, T. Nakajima, Y. Honda, O. Kitao, H. Nakai, T. Vreven, K. Throssell, J.A. Montgomery Jr., J.E. Peralta, F. Ogliaro, M.J. Bearpark, J.J. Heyd, E.N. Brothers, K.N. Kudin, V.N. Staroverov, T.A. Keith, R. Kobayashi, J. Normand, K. Raghavachari, A.P. Rendell, J.C. Burant, S.S. Iyengar, J. Tomasi, M. Cossi, J.M. Millam, M. Klene, C. Adamo, R. Cammi, J.W. Ochterski, R.L. Martin, K. Morokuma, O. Farkas, J.B. Foresman, D.J. Fox, Gaussian 16 Rev. A.03, (2016).

[31] J. Wang, W. Wang, P.A. Kollman, D.A. Case, Automatic atom type and bond type perception in molecular mechanical calculations, J. Mol. Graph. Model. 25 (2006) 247–260. https://doi.org/10.1016/j.jmgm.2005.12.005.

[32] J. Wang, R.M. Wolf, J.W. Caldwell, P.A. Kollman, D.A. Case, Development and testing of a general amber force field, J. Comput. Chem. 25 (2004) 1157–1174. https://doi.org/10.1002/jcc.20035.

[33] J.L. Perry, N. Dembla-Rajpal, L.A. Hall, J.B. Pritchard, A Three-dimensional Model of Human Organic Anion Transporter 1: AROMATIC AMINO ACIDS REQUIRED FOR SUBSTRATE TRANSPORT, J. Biol. Chem. 281 (2006) 38071–38079. https://doi.org/10.1074/jbc.M608834200.

[34] A.N. Rizwan, W. Krick, G. Burckhardt, The chloride dependence of the human organic anion transporter 1 (hOAT1) is blunted by mutation of a single amino acid, J. Biol. Chem. 282 (2007) 13402–13409. https://doi.org/10.1074/jbc.M609849200.

[35] L. Zou, A. Stecula, A. Gupta, B. Prasad, H.-C. Chien, S.W. Yee, L. Wang, J.D. Unadkat, S.H. Stahl, K.S. Fenner, K.M. Giacomini, Molecular Mechanisms for Species Differences in Organic Anion Transporter 1, OAT1: Implications for Renal Drug Toxicity, Mol. Pharmacol. 94 (2018) 689–699. https://doi.org/10.1124/mol.117.111153.

[36] J.A. Maier, C. Martinez, K. Kasavajhala, L. Wickstrom, K.E. Hauser, C. Simmerling, ff14SB: Improving the Accuracy of Protein Side Chain and Backbone Parameters from ff99SB, J. Chem. Theory Comput. 11 (2015) 3696–3713. https://doi.org/10.1021/acs.jctc.5b00255.

[37] C.J. Dickson, B.D. Madej, Å.A. Skjevik, R.M. Betz, K. Teigen, I.R. Gould, R.C. Walker, Lipid14: The Amber Lipid Force Field, J. Chem. Theory Comput. 10 (2014) 865–879. https://doi.org/10.1021/ct4010307.

[38] W.L. Jorgensen, J. Chandrasekhar, J.D. Madura, R.W. Impey, M.L. Klein, Comparison of simple potential functions for simulating liquid water, J. Chem. Phys. 79 (1983) 926–935. https://doi.org/10.1063/1.445869.

[39] I.S. Joung, T.E. Cheatham, Determination of Alkali and Halide Monovalent Ion Parameters for Use in Explicitly Solvated Biomolecular Simulations, J. Phys. Chem. B. 112 (2008) 9020–9041. https://doi.org/10.1021/jp8001614.

[40] I.S. Joung, T.E. Cheatham, Molecular Dynamics Simulations of the Dynamic and Energetic Properties of Alkali and Halide Ions Using Water-Model-Specific Ion Parameters, J. Phys. Chem. B. 113 (2009) 13279–13290. https://doi.org/10.1021/jp902584c.

[41] D.A. Case, I.Y. Ben-Shalom, S.R. Brozell, D.S. Cerutti, T.E. Cheatham, III, V.W.D. Cruzeiro, T.A. Darden, R.E. Duke, D. Ghoreishi, M.K. Gilson, H. Gohlke, A.W. Goetz, D. Greene, R Harris, N. Homeyer, Y. Huang, S. Izadi, A. Kovalenko, T. Kurtzman, T.S. Lee, S. LeGrand, P. Li, C. Lin, J. Liu, T. Luchko, R. Luo, D.J. Mermelstein, K.M. Merz, Y. Miao, G. Monard, C. Nguyen, H. Nguyen, I. Omelyan, A. Onufriev, F. Pan, R. Qi, D.R. Roe, A. Roitberg, C. Sagui, S. Schott-Verdugo, J. Shen, C.L. Simmerling, J. Smith, R. Salomon-Ferrer, J. Swails, R.C. Walker, J. Wang, H. Wei, R.M. Wolf, X. Wu, L. Xiao, D.M. York and P.A. Kollman, AMBER 2018, (2018).

[42] A.W. Götz, M.J. Williamson, D. Xu, D. Poole, S. Le Grand, R.C. Walker, Routine Microsecond Molecular Dynamics Simulations with AMBER on GPUs. 1. Generalized Born, J. Chem. Theory Comput. 8 (2012) 1542–1555. https://doi.org/10.1021/ct200909j.

[43] T. Darden, D. York, L. Pedersen, Particle mesh Ewald: An N ⋅log(N) method for Ewald sums in large systems, J. Chem. Phys. 98 (1993) 10089–10092. https://doi.org/10.1063/1.464397.

[44] R.J. Loncharich, B.R. Brooks, R.W. Pastor, Langevin dynamics of peptides: The frictional dependence of isomerization rates ofN-acetylalanyl-N?-methylamide, Biopolymers. 32 (1992) 523–535. https://doi.org/10.1002/bip.360320508.

[45] J. Åqvist, P. Wennerström, M. Nervall, S. Bjelic, B.O. Brandsdal, Molecular dynamics simulations of water and biomolecules with a Monte Carlo constant pressure algorithm, Chem. Phys. Lett. 384 (2004) 288–294. https://doi.org/10.1016/j.cplett.2003.12.039.

[46] D.R. Roe, T.E. Cheatham, PTRAJ and CPPTRAJ: Software for Processing and Analysis of Molecular Dynamics Trajectory Data, J. Chem. Theory Comput. 9 (2013) 3084–3095. https://doi.org/10.1021/ct400341p.

[47] W. Humphrey, A. Dalke, K. Schulten, VMD: Visual molecular dynamics, J. Mol. Graph. 14 (1996) 33–38. https://doi.org/10.1016/0263-7855(96)00018-5.

[48] A.M. Westerlund, O. Fleetwood, S. Pérez-Conesa, L. Delemotte, Network analysis reveals how lipids and other cofactors influence membrane protein allostery, J. Chem. Phys. 153 (2020) 141103. https://doi.org/10.1063/5.0020974.

[49] T. Cihlar, D.C. Lin, J.B. Pritchard, M.D. Fuller, D.B. Mendel, D.H. Sweet, The Antiviral Nucleotide Analogs Cidofovir and Adefovir Are Novel Substrates for Human and Rat Renal Organic Anion Transporter 1, Mol. Pharmacol. 56 (1999) 570–580. https://doi.org/10.1124/mol.56.3.570.

[50] T. Sekine, H. Miyazaki, H. Endou, Molecular physiology of renal organic anion transporters, Am. J. Physiol.-Ren. Physiol. 290 (2006) F251–F261. https://doi.org/10.1152/ajprenal.00439.2004.

[51] X.C. Zhang, Y. Zhao, J. Heng, D. Jiang, Energy coupling mechanisms of MFS transporters: Energy Coupling Mechanisms of MFS Transporters, Protein Sci. 24 (2015) 1560–1579. https://doi.org/10.1002/pro.2759.

[52] F. Di Meo, G. Fabre, K. Berka, T. Ossman, B. Chantemargue, M. Paloncýová, P. Marquet, M. Otyepka, P. Trouillas, In silico pharmacology: Drug membrane partitioning and crossing, Pharmacol. Res. 111 (2016) 471–486. https://doi.org/10.1016/j.phrs.2016.06.030.

[53] S. Li, Q. Zhang, G. You, Three ubiquitination sites of organic anion transporter-1 synergistically mediate protein kinase C-dependent endocytosis of the transporter, Mol. Pharmacol. 84 (2013) 139–146. https://doi.org/10.1124/mol.113.086769.

[54] M. Hong, F. Zhou, K. Lee, G. You, The putative transmembrane segment 7 of human organic anion transporter hOAT1 dictates transporter substrate binding and stability, J. Pharmacol. Exp. Ther. 320 (2007) 1209–1215. https://doi.org/10.1124/jpet.106.117663.

[55] J. Jumper, R. Evans, A. Pritzel, T. Green, M. Figurnov, O. Ronneberger, K. Tunyasuvunakool, R. Bates, A. Žídek, A. Potapenko, A. Bridgland, C. Meyer, S.A.A. Kohl, A.J. Ballard, A. Cowie, B. Romera-Paredes, S. Nikolov, R. Jain, J. Adler, T. Back, S. Petersen, D. Reiman, E. Clancy, M. Zielinski, M. Steinegger, M. Pacholska, T. Berghammer, S. Bodenstein, D. Silver, O. Vinyals, A.W. Senior, K. Kavukcuoglu, P. Kohli, D. Hassabis, Highly accurate protein structure prediction with AlphaFold, Nature. 596 (2021) 583–589. https://doi.org/10.1038/s41586-021-03819-2.

[56] V. Corradi, B.I. Sejdiu, H. Mesa-Galloso, H. Abdizadeh, S.Yu. Noskov, S.J. Marrink, D.P. Tieleman, Emerging Diversity in Lipid–Protein Interactions, Chem. Rev. 119 (2019) 5775–5848. https://doi.org/10.1021/acs.chemrev.8b00451.

[57] K. Kapoor, S. Pant, E. Tajkhorshid, Active Participation of Membrane Lipids in Inhibition of Multidrug Transporter P-Glycoprotein, BioRxiv. (2020) 2020.11.15.383794. https://doi.org/10.1101/2020.11.15.383794.

[58] D. Casares, P.V. Escribá, C.A. Rosselló, Membrane Lipid Composition: Effect on Membrane and Organelle Structure, Function and Compartmentalization and Therapeutic Avenues, Int. J. Mol. Sci. 20 (2019) 2167. https://doi.org/10.3390/ijms20092167.

[59] P.V. Escribá, X. Busquets, J. Inokuchi, G. Balogh, Z. Török, I. Horváth, J.L. Harwood, L. Vígh, Membrane lipid therapy: Modulation of the cell membrane composition and structure as a molecular base for drug discovery and new disease treatment, Prog. Lipid Res. 59 (2015) 38–53. https://doi.org/10.1016/j.plipres.2015.04.003.

[60] K. Bleasby, L.A. Hall, J.L. Perry, H.W. Mohrenweiser, J.B. Pritchard, Functional Consequences of Single Nucleotide Polymorphisms in the Human Organic Anion Transporter hOAT1 (SLC22A6), J. Pharmacol. Exp. Ther. 314 (2005) 923–931. https://doi.org/10.1124/jpet.105.084301.

[61] Z. Li, Current Updates in the Genetic Polymorphisms of Human Organic Anion Transporters (OATs), J. Pharmacogenomics Pharmacoproteomics. 03 (2012). https://doi.org/10.4172/2153-0645.1000e127.

[62] S.W. Yee, D.J. Brackman, E.A. Ennis, Y. Sugiyama, L.K. Kamdem, R. Blanchard, A. Galetin, L. Zhang, K.M. Giacomini, Influence of Transporter Polymorphisms on Drug Disposition and Response: A Perspective From the International Transporter Consortium, Clin. Pharmacol. Ther. 104 (2018) 803–817. https://doi.org/10.1002/cpt.1098.

[63] T. Fujita, C. Brown, E.J. Carlson, T. Taylor, M. de la Cruz, S.J. Johns, D. Stryke, M. Kawamoto, K. Fujita, R. Castro, C.-W. Chen, E.T. Lin, C.M. Brett, E.G. Burchard, T.E. Ferrin, C.C. Huang, M.K. Leabman, K.M. Giacomini, Functional analysis of polymorphisms in the organic anion transporter, SLC22A6 (OAT1):, Pharmacogenet. Genomics. 15 (2005) 201–209. https://doi.org/10.1097/01213011-200504000-00003.

[64] R.A. McPherson, M.R. Pincus, eds., Henry’s clinical diagnosis and management by laboratory methods, 23rd edition, Elsevier, St. Louis, Missouri, 2017.

[65] A. Emami Riedmaier, A.T. Nies, E. Schaeffeler, M. Schwab, Organic Anion Transporters and Their Implications in Pharmacotherapy, Pharmacol. Rev. 64 (2012) 421–449. https://doi.org/10.1124/pr.111.004614.

[66] G. Xu, V. Bhatnagar, G. Wen, B.A. Hamilton, S.A. Eraly, S.K. Nigam, Analyses of coding region polymorphisms in apical and basolateral human organic anion transporter (OAT) genes [OAT1 (NKT), OAT2, OAT3, OAT4, URAT (RST)], Kidney Int. 68 (2005) 1491–1499. https://doi.org/10.1111/j.1523-1755.2005.00612.x.

[67] I.M. da Rocha, A.S. Gasparotto, R.K. Lazzaretti, R.K. Notti, E. Sprinz, V.S. Mattevi, Polymorphisms associated with renal adverse effects of antiretroviral therapy in a Southern Brazilian HIV cohort, Pharmacogenet. Genomics. 25 (2015) 541–547. https://doi.org/10.1097/FPC.0000000000000169.

[68] J. Lowenstein, J.J. Grantham, The rebirth of interest in renal tubular function, Am. J. Physiol.-Ren. Physiol. 310 (2016) F1351–F1355. https://doi.org/10.1152/ajprenal.00055.2016.

[69] K.T. Bush, P. Singh, S.K. Nigam, Gut-derived uremic toxin handling in vivo requires OAT-mediated tubular secretion in chronic kidney disease, JCI Insight. 5 (2020) 133817. https://doi.org/10.1172/jci.insight.133817.

[70] T.T.G. Nieskens, J.G.P. Peters, M.J. Schreurs, N. Smits, R. Woestenenk, K. Jansen, T.K. van der Made, M. Röring, C. Hilgendorf, M.J. Wilmer, R. Masereeuw, A Human Renal Proximal Tubule Cell Line with Stable Organic Anion Transporter 1 and 3 Expression Predictive for Antiviral-Induced Toxicity, AAPS J. 18 (2016) 465–475. https://doi.org/10.1208/s12248-016-9871-8.

