## Supplementary material for "Substrate binding and lipid-mediated allostery in the human organic anion transporter 1 at the atomic-scale": Electronic Supplementary information

Electronic Supplementary Information for

INSERM U1248 Pharmacology & Transplantation, CBRS, Faculté de Médecine et Pharmacie, Univ. Limoges,

2 rue du Prof. Descottes,

87000 Limoges

France

### List of Tables

**Table S1.** Definitions of search volumes used during molecular docking calculations. Coordinates are centred on *h*OAT1 model center-of-mass. ... 3
**Table S2.** Total number of POPC ($N_{POPC}$), POPE ($N_{POPE}$), cholesterol ($N_{Chol}$), water ($N_{water}$) molecules and atoms ($N_{atoms}$) as well as initial PBC box sizes in MD simulations of (A) $\alpha$KG-bound *h*OAT1, (B) AFV-bound *h*OAT1 and (C) $\alpha$KG-AFV-bound *h*OAT1 simulations. ... 4
**Table S3.** Residues used in allostery pathway calculations, performed between intracellular domains and binding pockets ... 9
**Table S4.** Key H-bond network and contacts between adefovir and α-KG with *h*OAT1. H-bond fraction values greater than 1 represent multiple H-bonds between two residues (the maximum value can be 2.0 accounting for two H-bonds). ... 10
**Table S5.** *In vitro* studies of site-directed mutagenesis performed on OAT1. ... 11
**Table S6.** H-bond fractions between key residues of charge-relay system comparing apo and substrate-bound systems. The table considers simulations where $\alpha$KG is bound in (i) C-bundle triad, (ii) N-bundle triad and (iii) at the N/C-bundle interface for **(A)** $\alpha$KG-bound and **(B)** adefovir-$\alpha$KG-bound systems. H-bond fraction values greater than 1 represent multiple H-bonds between two residues (the maximum value can be 2.0 accounting for two H-bonds). ... 12

### List of Figures

**Figure S1.** The projection of initial snapshots of *h*OAT1 used for molecular docking and MD simulations with substrates onto the MFS conformational space resulted from PCA in POPC:POPE:Chol (2:1:1) (left), POPC:Chol (3:1) (middle) and POPC pure (right). ... 14
**Figure S2.** Molecular representations of search volumes used for molecular docking calculations. Search volumes for AFV (66560 Å$^3$) and a-KG (64768 Å$^3$) are depicted in blue and red, respectively. ... 15
**Figure S3.** Selected poses for adefovir (red sphere) and $\alpha$KG (green spheres) from molecular docking calculations. ... 16
**Figure S4.** Time-dependent root mean square deviation (RMSD) of substrate-bound simulations in different lipid bilayer membranes, namely POPC:POPE:Chol (2:1:1), POPC:Chol (3:1) and POPC. First frame was used as reference frame. ... 17
**Figure S5.** Efficiencies of the allosteric pathways from *h*OAT1 A-motifs to (a) inner binding pocket and (b) B-like motif. For each panel, N- and C-bundle efficiencies are provided in top and bottom, respectively. Calculations were performed considering POPC:POPE:Chol (2:1:1) lipid bilayer. ... 18
**Figure S6.** Efficiencies of the allosteric pathways from *h*OAT1 E[$X_6$]R motifs to (a) inner binding pocket and (b) B-like motif. For each panel, N- and C-bundle efficiencies are provided in top and bottom, respectively. Calculations were performed considering POPC:POPE:Chol (2:1:1) lipid bilayer. ... 19
**Figure S7.** Efficiencies of the allosteric pathways from *h*OAT1 (a) inner binding pocket and (b) b-like motif to A-motifs. For each panel, N- and C-bundle efficiencies are provided in top and bottom, respectively. Calculations were performed considering POPC:POPE:Chol (2:1:1) lipid bilayer. ... 20
**Figure S8.** Efficiencies of the allosteric pathways from *h*OAT1 (a) inner binding pocket and (b) b-like motif to E[$X_6$]R-motifs. For each panel, N- and C-bundle efficiencies are provided in top and bottom, respectively. Calculations were performed in POPC:POPE:Chol (2:1:1) lipid bilayer. ... 21
**Figure S9.** Calculated *h*OAT1 per-residue involvement (betweenness) in the allosteric pathway from B-like motif to N-bundle (left) and C-bundle (right) A-motifs for (a) apo, (b) adefovir-bound, (c) $\alpha$KG-bound and (d) adefovir-$\alpha$KG-bound *h*OAT1 systems in POPC:POPE:Chol (2:1:1). TMH topologies are depicted in light blue background. ... 22
**Figure S10.** Calculated *h*OAT1 per-residue involvement (betweenness) in the allosteric pathway from inner binding pocket to N-bundle (left) and C-bundle (right) A-motifs for (a) apo, (b) adefovir-bound, (c) $\alpha$KG-bound and (d) adefovir-$\alpha$KG-bound *h*OAT1 systems in POPC:POPE:Chol (2:1:1). TMH topologies are depicted in light blue background. ... 23
**Figure S11.** Calculated *h*OAT1 per-residue involvement (betweenness) in the allosteric pathway from B-like motif to N-bundle (left) and C-bundle (right) E[$X_6$]R for (a) apo, (b) adefovir-bound, (c) $\alpha$KG-bound and (d) adefovir-$\alpha$KG-bound *h*OAT1 systems. TMH topologies are depicted in light blue background. ... 24
**Figure S12.** Calculated *h*OAT1 per-residue involvement (betweenness) in the allosteric pathway from inner binding cavity to N-bundle (left) and C-bundle (right) E[$X_6$]R for (a) apo, (b) adefovir-bound, (c) $\alpha$KG-bound and (d) adefovir-$\alpha$KG-bound *h*OAT1 systems. TMH topologies are depicted in light blue background. ... 25
**Figure S13.** Structures of adefovir, tenofovir, cidofovir, ochratoxin A, p-aminohippuric acid, methotrexate. ... 26

### 1. Supplemental Tables

**Table S1.** Definitions of search volumes used during molecular docking calculations. Coordinates are centred on *h*OAT1 model center-of-mass.

| **Volume search space** | **Box center (Å)** | | | **Box dimensions (Å)** | | | **Substrate** |
|---|---|---|---|---|---|---|---|
| | *x* | *y* | *z* | *x* | *y* | *z* | |
| Extracellular side | 40 | 156 | 79 | 52.0 | 40.0 | 32.0 | AFV |
| Intracellular side | 43 | 155 | 48 | 46.0 | 44.0 | 32.0 | $\alpha$KG |

**Table S2.** Total number of POPC ($N_{POPC}$), POPE ($N_{POPE}$), cholesterol ($N_{Chol}$), water ($N_{water}$) molecules and atoms ($N_{atoms}$) as well as initial PBC box sizes in MD simulations of (A) $\alpha$KG-bound *h*OAT1, (B) AFV-bound *h*OAT1 and (C) $\alpha$KG-AFV-bound *h*OAT1 simulations.

**(A) $\alpha$KG-bound *h*OAT1**

*Position 1*

| **POPC:POPE:Chol ratio** | | | **1:0:0** | **3:0:1** | **2:1:1** |
|---|---|---|---|---|---|
| Lipids | $N_{POPC}$ | Upper leaflet | 189 | 156 | 108 |
| | | Lower leaflet | 184 | 159 | 108 |
| | | Total | 373 | 315 | 216 |
| | $N_{POPE}$ | Upper leaflet | - | - | 54 |
| | | Lower leaflet | - | - | 54 |
| | | Total | - | - | 108 |
| | $N_{chol}$ | Upper leaflet | - | 52 | 54 |
| | | Lower leaflet | - | 53 | 54 |
| | | Total | - | 105 | 108 |
| Water | $N_{water}$ | | 39384 | 44522 | 42570 |
| Box size | x | | 118 | 113 | 113 |
| (Å) | y | | 117 | 114 | 113 |
| | z | | 125 | 145 | 141 |
| $N_{atoms}$ | | | 176919 | 192363 | 186949 |

*Position 2*

| **POPC:POPE:Chol ratio** | | | **1:0:0** | **3:0:1** | **2:1:1** |
|---|---|---|---|---|---|
| Lipids | $N_{POPC}$ | Upper leaflet | 187 | 159 | 110 |
| | | Lower leaflet | 186 | 159 | 110 |
| | | Total | 373 | 318 | 220 |
| | $N_{POPE}$ | Upper leaflet | - | - | 55 |
| | | Lower leaflet | - | - | 55 |
| | | Total | - | - | 110 |
| | $N_{chol}$ | Upper leaflet | - | 53 | 55 |
| | | Lower leaflet | - | 53 | 55 |
| | | Total | - | 106 | 110 |
| Water | $N_{water}$ | | 40301 | 42007 | 43089 |
| Box size | x | | 120 | 115 | 115 |
| (Å) | y | | 120 | 112 | 113 |
| | z | | 122 | 139 | 141 |
| Natoms | | | 179672 | 189442 | 189442 |

*Position 3*

| POPC:POPE:Chol ratio | | | 1:0:0 | 3:0:1 | 2:1:1 |
|---|---|---|---|---|---|
| Lipids | $N_{POPC}$ | Upper leaflet | 185 | 159 | 110 |
| | | Lower leaflet | 186 | 159 | 110 |
| | | Total | 371 | 318 | 220 |
| | $N_{POPE}$ | Upper leaflet | - | - | 55 |
| | | Lower leaflet | - | - | 55 |
| | | Total | - | - | 110 |
| | $N_{chol}$ | Upper leaflet | - | 53 | 55 |
| | | Lower leaflet | - | 53 | 55 |
| | | Total | - | 106 | 110 |
| Water | $N_{water}$ | | 41576 | 41247 | 40538 |
| Box size (Å) | x | | 118 | 113 | 114 |
| | y | | 118 | 114 | 113 |
| | z | | 129 | 138 | 142 |
| Natoms | | | 183235 | 182856 | 181773 |

**(B) AFV-bound *h*OAT1**

*Position 1*

| POPC:POPE:Chol ratio | | | 1:0:0 | 3:0:1 | 2:1:1 |
|---|---|---|---|---|---|
| Lipids | $N_{POPC}$ | Upper leaflet | 189 | 156 | 108 |
| | | Lower leaflet | 184 | 159 | 108 |
| | | Total | 373 | 315 | 216 |
| | $N_{POPE}$ | Upper leaflet | - | - | 54 |
| | | Lower leaflet | - | - | 54 |
| | | Total | - | - | 108 |
| | $N_{chol}$ | Upper leaflet | - | 52 | 54 |
| | | Lower leaflet | - | 53 | 54 |
| | | Total | - | 105 | 108 |
| Water | $N_{water}$ | | 39361 | 44449 | 42550 |
| Box size (Å) | x | | 119 | 112 | 115 |
| | y | | 119 | 113 | 113 |
| | z | | 122 | 147 | 140 |
| Natoms | | | 176850 | 192144 | 186891 |

*Position 2*

| POPC:POPE:Chol ratio | | | 1:0:0 | 3:0:1 | 2:1:1 |
|---|---|---|---|---|---|
| Lipids | $N_{POPC}$ | Upper leaflet | 187 | 159 | 110 |
| | | Lower leaflet | 189 | 159 | 110 |
| | | Total | 373 | 318 | 220 |
| | $N_{POPE}$ | Upper leaflet | - | - | 55 |
| | | Lower leaflet | - | - | 55 |
| | | Total | - | - | 110 |
| | $N_{chol}$ | Upper leaflet | - | 53 | 55 |
| | | Lower leaflet | - | 53 | 55 |
| | | Total | - | 106 | 110 |
| Water | $N_{water}$ | | 40284 | 42053 | 43119 |
| Box size | x | | 118 | 112 | 113 |
| (Å) | y | | 118 | 112 | 113 |
| | z | | 125 | 142 | 143 |
| Natoms | | | 179621 | 185416 | 189532 |

*Position 3*

| POPC:POPE:Chol ratio | | | 1:0:0 | 3:0:1 | 2:1:1 |
|---|---|---|---|---|---|
| Lipids | $N_{POPC}$ | Upper leaflet | 185 | 159 | 110 |
| | | Lower leaflet | 186 | 159 | 110 |
| | | Total | 371 | 318 | 220 |
| | $N_{POPE}$ | Upper leaflet | - | - | 55 |
| | | Lower leaflet | - | - | 55 |
| | | Total | - | - | 110 |
| | $N_{chol}$ | Upper leaflet | - | 53 | 55 |
| | | Lower leaflet | - | 53 | 55 |
| | | Total | - | 106 | 110 |
| Water | $N_{water}$ | | 41587 | 41252 | 40556 |
| Box size | x | | 118 | 114 | 115 |
| (Å) | y | | 118 | 113 | 112 |
| | z | | 128 | 138 | 136 |
| Natoms | | | 183270 | 182856 | 181829 |

**(C) $\alpha$KG-AFV-bound *h*OAT1**

*Position 1*

| **POPC:POPE:Chol ratio** | | | **1:0:0** | **3:0:1** | **2:1:1** |
|---|---|---|---|---|---|
| Lipids | $N_{POPC}$ | Upper leaflet | 189 | 156 | 108 |
| | | Lower leaflet | 184 | 159 | 108 |
| | | Total | 373 | 315 | 216 |
| | $N_{POPE}$ | Upper leaflet | - | - | 54 |
| | | Lower leaflet | - | - | 54 |
| | | Total | - | - | 108 |
| | $N_{chol}$ | Upper leaflet | - | 52 | 54 |
| | | Lower leaflet | - | 53 | 54 |
| | | Total | - | 105 | 108 |
| Water | $N_{water}$ | | 39383 | 44522 | 42570 |
| Box size | x | | 118 | 112 | 113 |
| (Å) | y | | 118 | 113 | 114 |
| | z | | 123 | 147 | 141 |
| Natoms | | | 176947 | 192391 | 186977 |

*Position 2*

| **POPC:POPE:Chol ratio** | | | **1:0:0** | **3:0:1** | **2:1:1** |
|---|---|---|---|---|---|
| Lipids | $N_{POPC}$ | Upper leaflet | 187 | 159 | 110 |
| | | Lower leaflet | 186 | 159 | 110 |
| | | Total | 373 | 318 | 220 |
| | $N_{POPE}$ | Upper leaflet | - | - | 55 |
| | | Lower leaflet | - | - | 55 |
| | | Total | - | - | 110 |
| | $N_{chol}$ | Upper leaflet | - | 53 | 55 |
| | | Lower leaflet | - | 53 | 55 |
| | | Total | - | 106 | 110 |
| Water | $N_{water}$ | | 40301 | 42007 | 43089 |
| Box size | x | | 120 | 115 | 114 |
| (Å) | y | | 120 | 112 | 112 |
| | z | | 122 | 140 | 144 |
| Natoms | | | 179672 | 185304 | 189470 |

*Position 3*

| **POPC:POPE:Chol ratio** | | | **1:0:0** | **3:0:1** | **2:1:1** |
|---|---|---|---|---|---|
| Lipids | $N_{POPC}$ | Upper leaflet | 185 | 159 | 110 |
| | | Lower leaflet | 186 | 159 | 110 |
| | | Total | 371 | 318 | 220 |
| | $N_{POPE}$ | Upper leaflet | - | - | 55 |
| | | Lower leaflet | - | - | 55 |
| | | Total | - | - | 110 |
| | $N_{chol}$ | Upper leaflet | - | 53 | 55 |
| | | Lower leaflet | - | 53 | 55 |
| | | Total | - | 106 | 110 |
| Water | $N_{water}$ | | 41575 | 41247 | 40538 |
| Box size | x | | 117 | 113 | 114 |
| (Å) | y | | 117 | 115 | 113 |
| | z | | 130 | 137 | 136 |
| Natoms | | | 183263 | 182884 | 181801 |

**Table S3.** Residues used in allostery pathway calculations, performed between intracellular domains and binding pockets.

| Motifs | Residues |
| --- | --- |
| B-like motif | Arg192 Leu193 Leu194 Ser195 Gly196 |
| The inner binding cavity | Arg466 Tyr230 Thr36 Ser350 Thr349 |
| A-motif (C-bundle) | Asn390 Ser391 Leu392 Gly393 Arg394 Arg395 |
| A-motif (N-bundle) | Asp157 Arg158 Leu159 Gly160 Arg161 Arg162 |
| E[$X_6$]R (N-bundle) | Glu212 Trp213 Met214 Pro215 Ile216 Ile217 Thr218 Arg219 |
| E[$X_6$]R (C-bundle) | Glu447 Leu448 Tyr449 Pro450 Thr451 Met452 Ile453 Arg454 |

**Table S4.** Key H-bond network and contacts between adefovir and α-KG with *h*OAT1. H-bond fraction values greater than 1 represent multiple H-bonds between two residues (the maximum value can be 2.0 accounting for two H-bonds).

| Substrate | Binding pocket | H-bond | | Contacts | |
|---|---|---|---|---|---|
| | | Residue | Fraction | Residue | Fraction |
| Adefovir | B-like motif cavity | Arg192 | 1.950 | Phe258 | 0.96 |
| | | Ser255 | 0.907 | Tyr141* | 0.94 |
| | | Gln251 | 0.573 | His34 | 0.92 |
| | | Ser195 | 0.504 | Gly196 | 0.78 |
| | | Tyr141 | 0.473 | Leu199 | 0.72 |
| | | Gly62 | 0.110 | Leu37 | 0.67 |
| | Inner cavity | Arg466 | 1.569 | Trp346 | 0.99 |
| | | Asn35 | 0.679 | Asn39 | 0.96 |
| | | Trp346 | 0.657 | Met142 | 0.89 |
| | | Ser469 | 0.600 | Ser350 | 0.88 |
| | | Gln138 | 0.461 | Tyr230 | 0.70 |
| | | Tyr230 | 0.377 | Gly468 | 0.68 |
| | | Gln234 | 0.112 | Ala32 | 0.60 |
| | | | | Met31 | 0.49 |
| $\alpha$KG | E[$X_6$]R Interface | Thr451 | 1.857 | Pro450 | 0.987 |
| | | Arg219 | 1.673 | Thr515 | 0.775 |
| | | Arg454 | 1.025 | Val516 | 0.540 |
| | | Arg522 | 0.593 | Arg522 | 0.498 |
| | | Arg273 | 0.403 | Hie217 | 0.488 |
| | | | | Val211 | 0.480 |
| | | | | Leu512 | 0.457 |
| | C-bundle triad | Arg219 | 1.515 | Leu443 | 0.892 |
| | | Arg394 | 1.376 | Ala220 | 0.824 |
| | | Arg454 | 1.008 | Cys440 | 0.659 |
| | | | | Gly223 | 0.588 |
| | | | | Ile389 | 0.235 |
| | N-bundle triad | Arg162 | 2.019 | Leu209 | 0.974 |
| | | Arg161 | 1.698 | Trp213 | 0.909 |
| | | Asn205 | 0.868 | Glu212 | 0.682 |
| | | Thr208 | 0.866 | Ser271 | 0.661 |
| | | Gln455 | 0.629 | | |

**Table S5.** *In vitro* studies of site-directed mutagenesis performed on OAT1.

| Mutation | Localization | Substrate | Expression | Transport | Cell model | Ref. |
|---|---|---|---|---|---|---|
| p.Leu30Ala | TMH1 | PAH | ↓ | ↓ | COS-7 | [1] |
| p.Thr36Ala<br>p.Thr36Ser<br>p.Thr36Cys | TMH1 | PAH | No effect | ↓ | COS-7 | |
| p.Asn39Glu | TMH1 | PAH | No effect | ↓ | HeLa | [2] |
| p.Asn56Glu | ECL1 | PAH | No effect | No effect | HeLa | |
| p.Asn92Glu | ECL1 | PAH | No effect | No effect | HeLa | |
| p.Asn97Glu | ECL1 | PAH | No effect | No effect | HeLa | |
| p.Asn113Glu | ECL1 | PAH | No effect | No effect | HeLa | |
| p.Ser203Ala | TMH4 | AFV, CDF, TNF | No data | ↓ | HEK293 | [3] |
| p.Tyr230Ala<br>p.Tyr230Phe | TMH5 | PAH | ↓ | ↓ | Xenopus Oocyte | [4] |
| p.Trp346Ala | TMH7 | None | ↓ | No data | COS7 | [5] |
| p.Thr349Ala | TMH7 | None | ↓ | No data | COS7 | |
| p.Tyr353Ala<br>p.Tyr353Trp<br>p.Tyr353Phe | TMH7 | PAH | No effect | ↓ | COS7 | |
| p.Tyr354Ala<br>p.Tyr354Trp<br>p.Tyr354Phe | TMH7 | PAH | No effect | ↓ | COS7 | |
| p.Lys431Ala<br>p.Lys431Arg | TMH10 | CDF, PAH | ↓ | ↓ | Xenopus Oocyte | [4] |
| p.Phe438Ala<br>p.Phe438Tyr | TMH10 | CDF, PAH | ↓ | ↓ | Xenopus Oocyte | |
| p.Arg466Lys<br>p.Arg466Asp | TMH11 | PAH | No effect | ↓ | Xenopus Oocyte | [6] |
| p.Tyr490Ala | TMH12 | PAH | ↓ | ↓ | COS-7 & LLC-PK1 | [7] |
| p.Tyr490Phe | | | No effect | No effect | | |
| p.Tyr490Tyr | | | ↓ | ↓ | | |
| p.Leu503Ala | TMH12 | PAH | ↓ | ↓ | COS-7 & LLC-PK1 | |
| p.Leu504Ala | TMH12 | PAH | ↓ | ↓ | COS-7 & LLC-PK1 | |
| p.Glu506Ala | ICH6 | PAH | No effect | ↓ | COS-7 | [8] |
| p.Glu506Gln | | | No effect | ↓ | | |
| p.Glu506Asp | | | No effect | No effect | | |
| p.Leu512Ala | ICH6 | PAH | No effect | ↓ | COS-7 | |
| p.Leu512Val | | | No effect | ↓ | | |
| p.Leu512Ile | | | No effect | No effect | | |

**Abbreviations:** AFV: adefovir disproxyl fumarate; CDF: cidofovir; PAH: *p*-aminohippurate; TNF, tenofovir; ↓ decrease.

**Table S6.** H-bond fractions between key residues of charge-relay system comparing apo and substrate-bound systems. The table considers simulations where $\alpha$KG is bound in (i) C-bundle triad, (ii) N-bundle triad and (iii) at the N/C-bundle interface for **(A)** $\alpha$KG-bound and **(B)** adefovir-$\alpha$KG-bound systems. H-bond fraction values greater than 1 represent multiple H-bonds between two residues (the maximum value can be 2.0 accounting for two H-bonds in "separately" columns).

(A)

| Domains | Motifs/TMHs | | H-bond | separately | | | | cumulative | | | |
|---|---|---|---|---|---|---|---|---|---|---|---|
| | | | | Apo | $\alpha$KG position: C-bundle triad | $\alpha$KG position: N-bundle triad | $\alpha$KG position: N-, C-bundle interface | Apo | $\alpha$KG position: N-bundle triad | $\alpha$KG position: C-bundle triad | $\alpha$KG position: N-, C-bundle interface |
| N-bundle | A-motif | PETL | Arg162-Glu270 | 0.569 | 0.471 | 0.123 | 0.267 | 4.102 | 4.111 | 2.288 | 3.896 |
| | | | Arg161-Ile269 | 0.126 | 1.112 | 0.000 | 0.000 | | | | |
| | A-motif | E[X$_6$]R | Arg161-Glu270 | 0.109 | 0.603 | 0.000 | 0.000 | | | | |
| | | | Arg161-Glu212 | 1.518 | 0.608 | 0.000 | 1.946 | | | | |
| | E[X$_6$]R | PETL | Glu212-Arg273 | 1.160 | 0.624 | 1.028 | 0.711 | | | | |
| | | | Glu212-Ser271 | 0.620 | 0.693 | 0.681 | 0.972 | | | | |
| | | | Trp213-Ile269 | 0.000 | 0.000 | 0.456 | 0.000 | | | | |
| C-bundle | A-motif | PETL | Arg394-Glu506 | 0.674 | 0.446 | 0.421 | 0.882 | 2.755 | 2.475 | 2.036 | 4.416 |
| | | | Arg395-Glu506 | 0.322 | 0.108 | 0.294 | 0.690 | | | | |
| | A-motif | E[X$_6$] | Arg394-Glu447 | 1.129 | 1.286 | 1.321 | 1.893 | | | | |
| | E[X$_6$]R | PETL | Glu447-Tyr507 | 0.630 | 0.635 | 0.000 | 0.951 | | | | |
| N- and C-bundle interface | E[X$_6$]R | E[X$_6$] | Arg219-Glu447 | 1.334 | 1.311 | 0.950 | 0.328 | 3.139 | 1.550 | 1.435 | 0.328 |
| | E[X$_6$]R | TMH11 | Glu212-Gln455 | 0.464 | 0.000 | 0.000 | 0.000 | | | | |
| | A-motif | TMH11 | Asp157-Gln455 | 0.224 | 0.000 | 0.000 | 0.000 | | | | |
| | TMH5 | A-motif | Thr224-Asn390 | 0.776 | 0.239 | 0.485 | 0.000 | | | | |
| | | | Ala220-Asn390 | 0.134 | 0.000 | 0.000 | 0.000 | | | | |

| | | | | | | | |
|---|---|---|---|---|---|---|---|
| | A-motif | TMH4 | Arg161-Asn205 | 0.207 | 0.000 | 0.000 | 0.000 |

(B)

| | | | | separately | | | | cumulative | | | |
|---|---|---|---|---|---|---|---|---|---|---|---|
| | | | | | $\alpha$KG position | | | | $\alpha$KG position | | |
| Domains | Motifs/TMHs | | H-bond | Apo | C-bundle triad | N-bundle triad | N-, C-bundle interface | Apo | N-bundle triad | C-bundle triad | N-, C-bundle interface |
| N-bundle | A-motif | PETL | Arg162-Glu270 | 0.569 | 0.638 | 0.531 | 0.364 | 4.102 | 4.370 | 3.338 | 3.622 |
| | | | Arg161-Ile269 | 0.126 | 0.225 | 0.000 | 0.000 | | | | |
| | A-motif | E[$X_6$]R | Arg161-Glu270 | 0.109 | 0.160 | 0.000 | 0.123 | | | | |
| | | | Arg161-Glu212 | 1.518 | 1.539 | 1.530 | 1.801 | | | | |
| | E[$X_6$]R | PETL | Glu212-Arg273 | 1.160 | 1.157 | 0.239 | 0.394 | | | | |
| | | | Glu212-Ser271 | 0.620 | 0.651 | 0.636 | 0.940 | | | | |
| | | | Trp213-Ile269 | 0.000 | 0.000 | 0.402 | 0.000 | | | | |
| C-bundle | A-motif | PETL | Arg394-Glu506 | 0.674 | 0.474 | 0.717 | 0.593 | 2.755 | 3.830 | 0.997 | 3.429 |
| | | | Arg395-Glu506 | 0.322 | 0.717 | 0.280 | 0.456 | | | | |
| | A-motif | E[$X_6$]R | Arg-394-Glu447 | 1.129 | 1.798 | 0.000 | 1.601 | | | | |
| | E[$X_6$]R | PETL | Glu447-Tyr507 | 0.630 | 0.841 | 0.000 | 0.779 | | | | |
| N- and C-bundle interface | E[$X_6$]R | E[$X_6$]R | Arg219-Glu447 | 1.334 | 0.000 | 1.056 | 0.805 | 3.139 | 0.520 | 2.087 | 1.720 |
| | E[$X_6$]R | TMH11 | Glu212-Gln455 | 0.464 | 0.000 | 0.235 | 0.000 | | | | |
| | A-motif | TMH11 | Asp157-Gln455 | 0.224 | 0.000 | 0.000 | 0.358 | | | | |
| | TMH5 | A-motif | Thr224-Asn390 | 0.776 | 0.000 | 0.391 | 0.315 | | | | |
| | | | Ala220-Asn390 | 0.134 | 0.000 | 0.000 | 0.000 | | | | |
| | A-motif | TMH4 | Arg161-Asn205 | 0.207 | 0.520 | 0.405 | 0.242 | | | | |

### 2. Supplemental figures

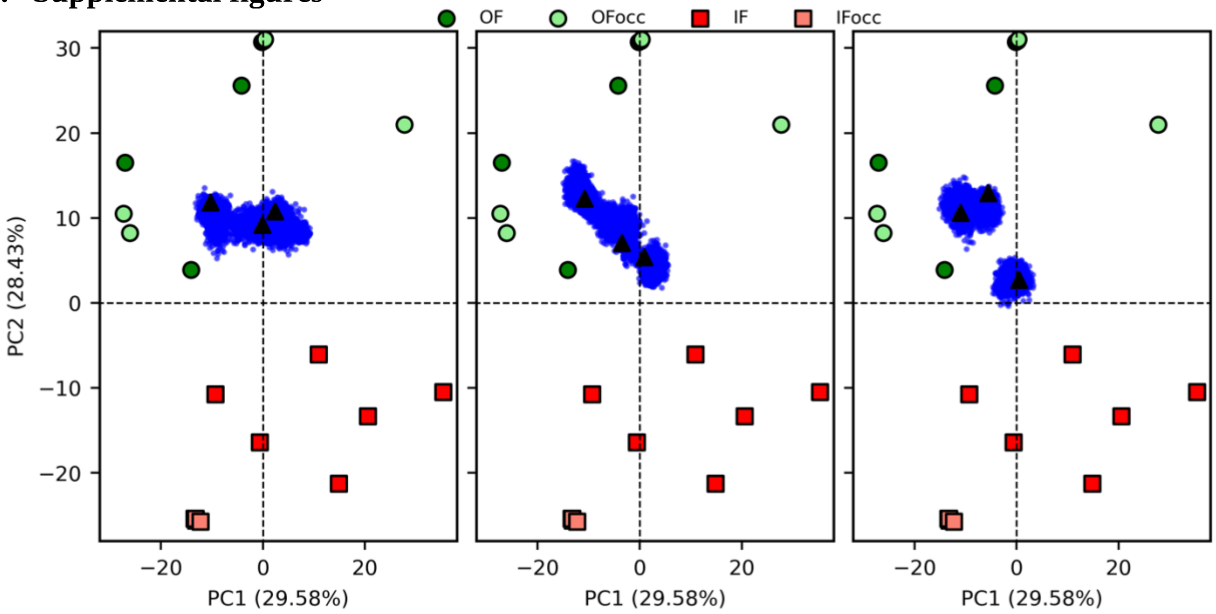


**Figure S1.** The projection of initial snapshots of *h*OAT1 used for molecular docking and MD simulations with substrates onto the MFS conformational space resulted from PCA in POPC:POPE:Chol (2:1:1) (left), POPC:Chol (3:1) (middle) and POPC pure (right).

Blue dots represent snaphots obtained from apo hOAT1 MD simulations and black triangle are the selected snapshots for molecular docking calculations. Description of apo hOAT1 MD simulations and analyses performed to create the present MFS conformational space are available in Ref. [9].

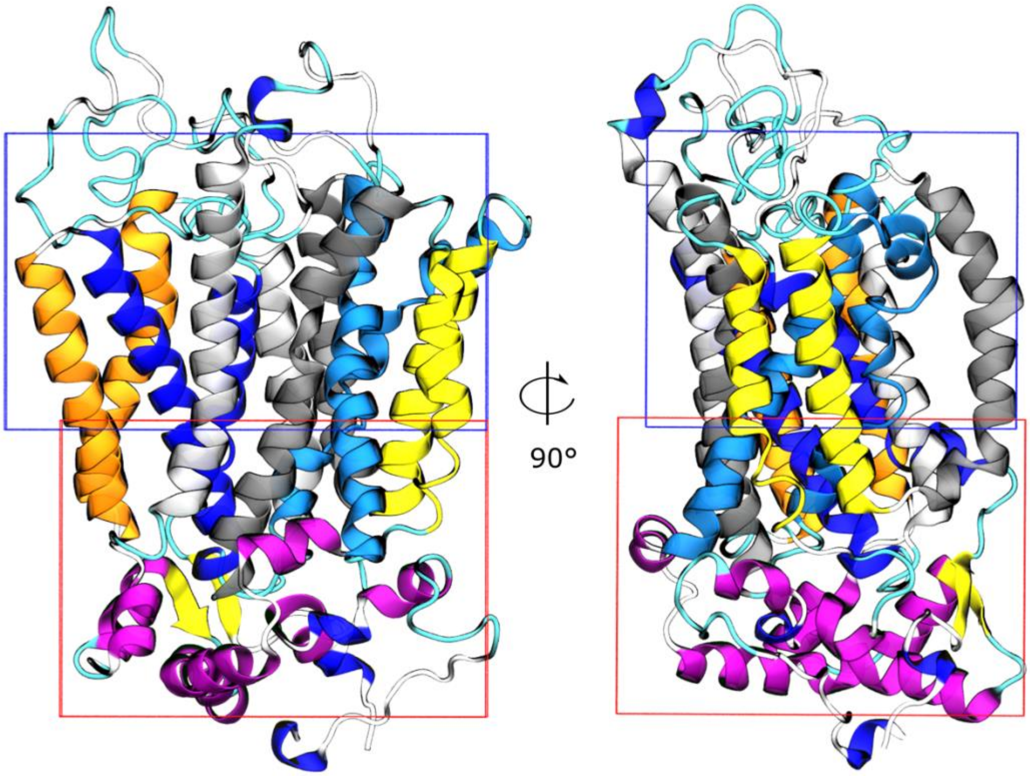


**Figure S2.** Molecular representations of search volumes used for molecular docking calculations. Search volumes for AFV (66560 Å$^3$) and a-KG (64768 Å$^3$) are depicted in blue and red, respectively.

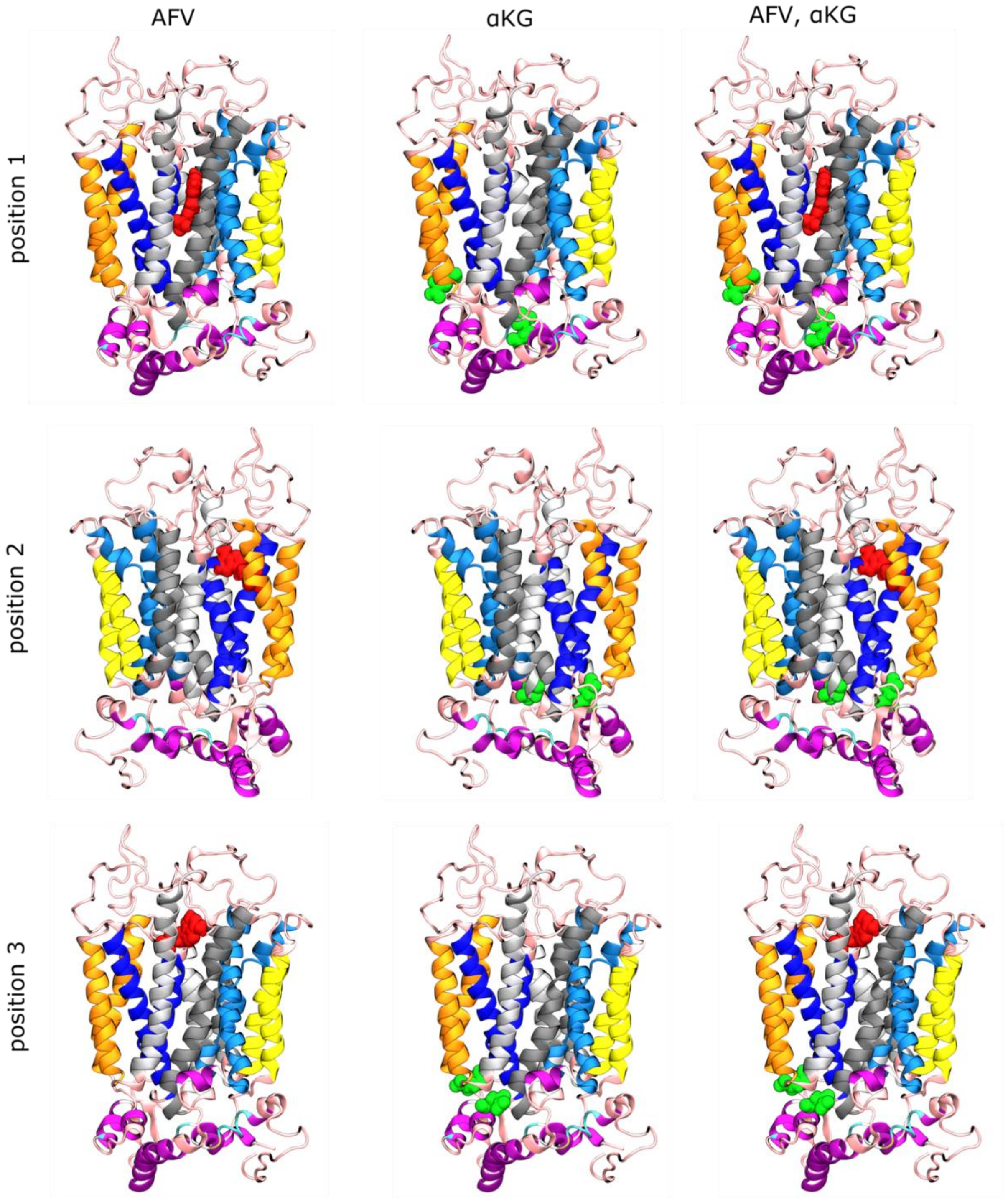


**Figure S3.** Selected poses for adefovir (red sphere) and $\alpha$KG (green spheres) from molecular docking calculations.

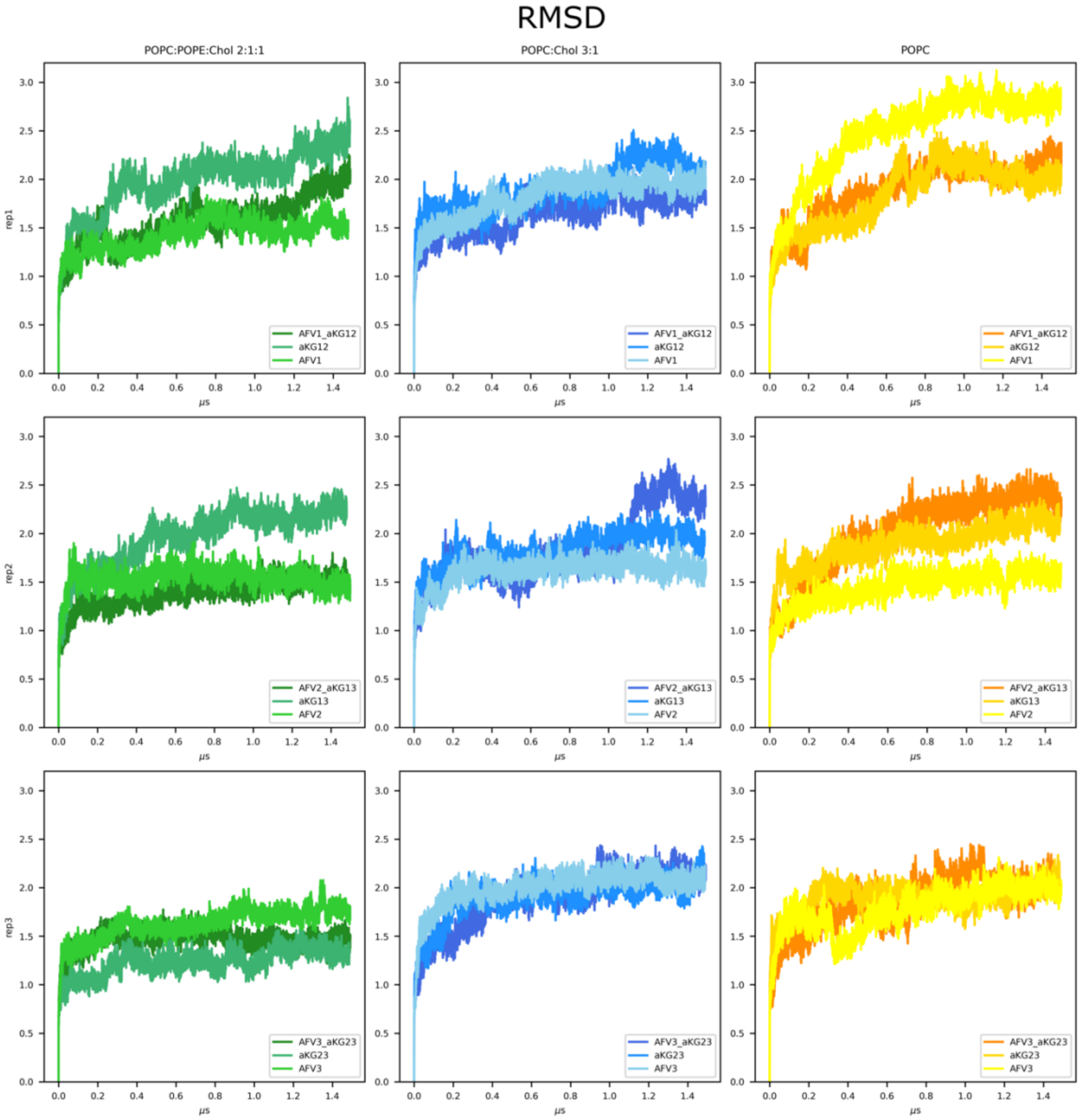


**Figure S4.** Time-dependent root mean square deviation (RMSD) of substrate-bound simulations in different lipid bilayer membranes, namely POPC:POPE:Chol (2:1:1), POPC:Chol (3:1) and POPC. First frame was used as reference frame.

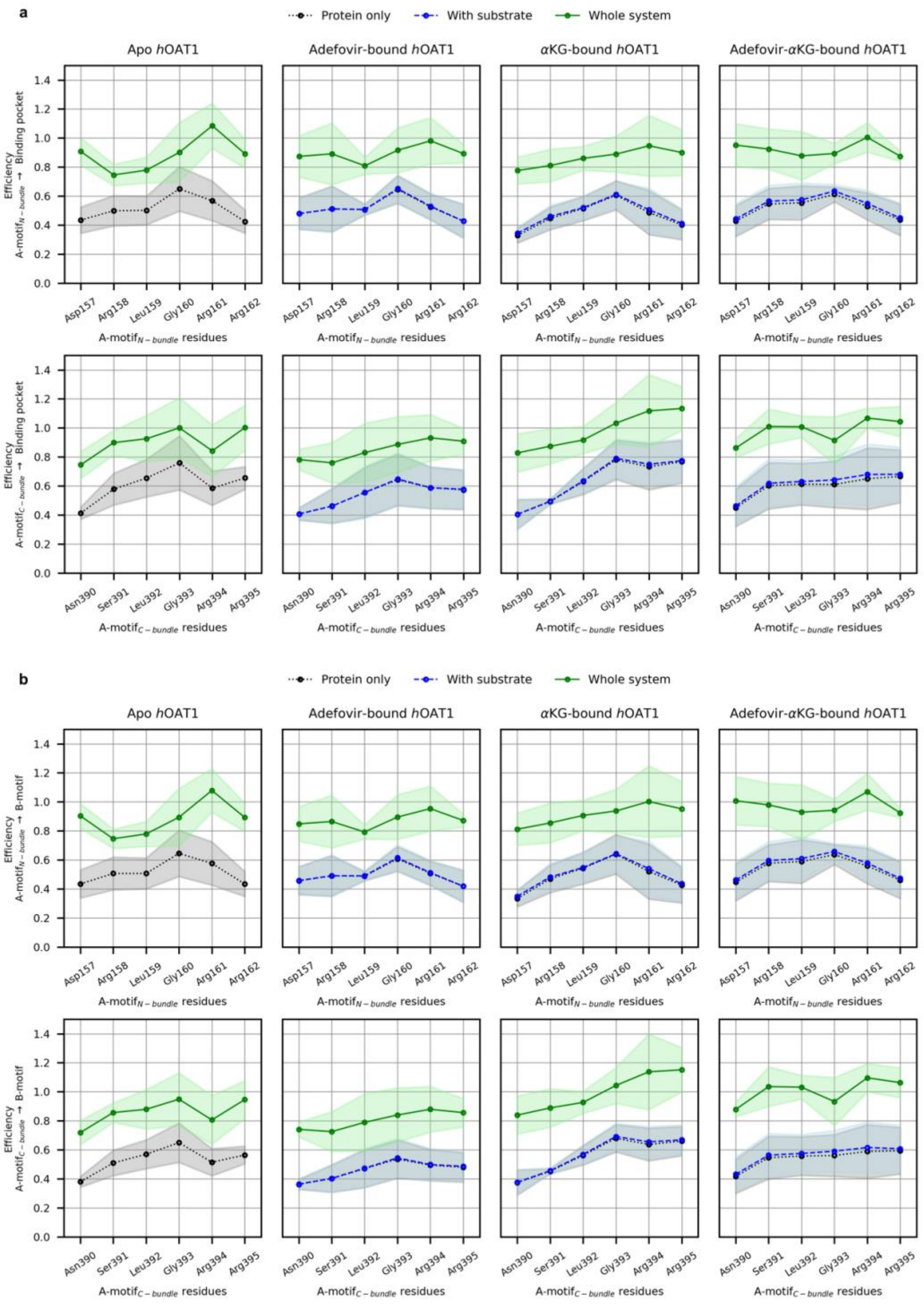


**Figure S5.** Efficiencies of the allosteric pathways from *h*OAT1 A-motifs to (a) inner binding pocket and (b) B-like motif. For each panel, N- and C-bundle efficiencies are provided in top and bottom, respectively. Calculations were performed considering POPC:POPE:Chol (2:1:1) lipid bilayer.

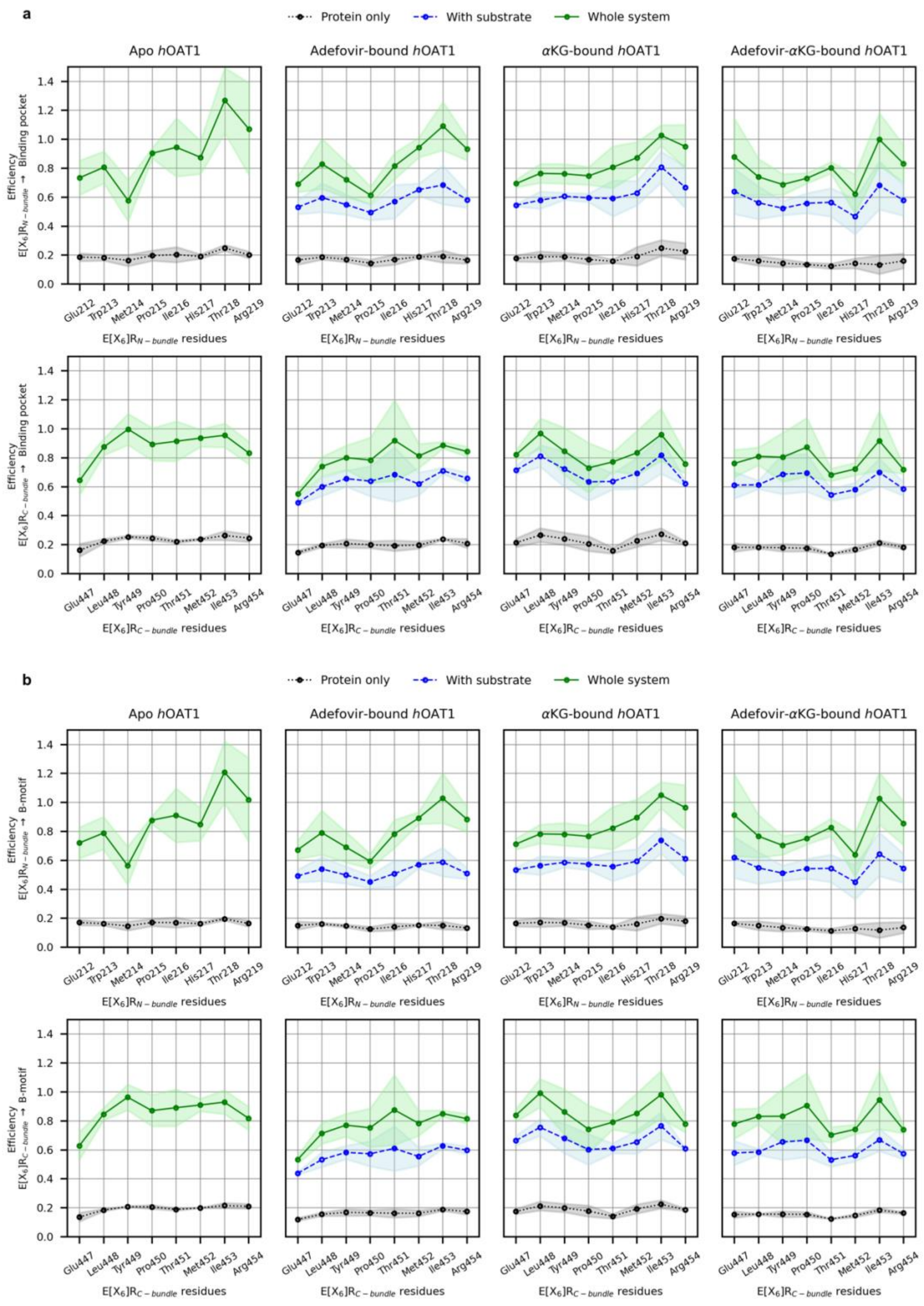


**Figure S6.** Efficiencies of the allosteric pathways from *h*OAT1 E[$X_6$]R motifs to (a) inner binding pocket and (b) B-like motif. For each panel, N- and C-bundle efficiencies are provided in top and bottom, respectively. Calculations were performed considering POPC:POPE:Chol (2:1:1) lipid bilayer.

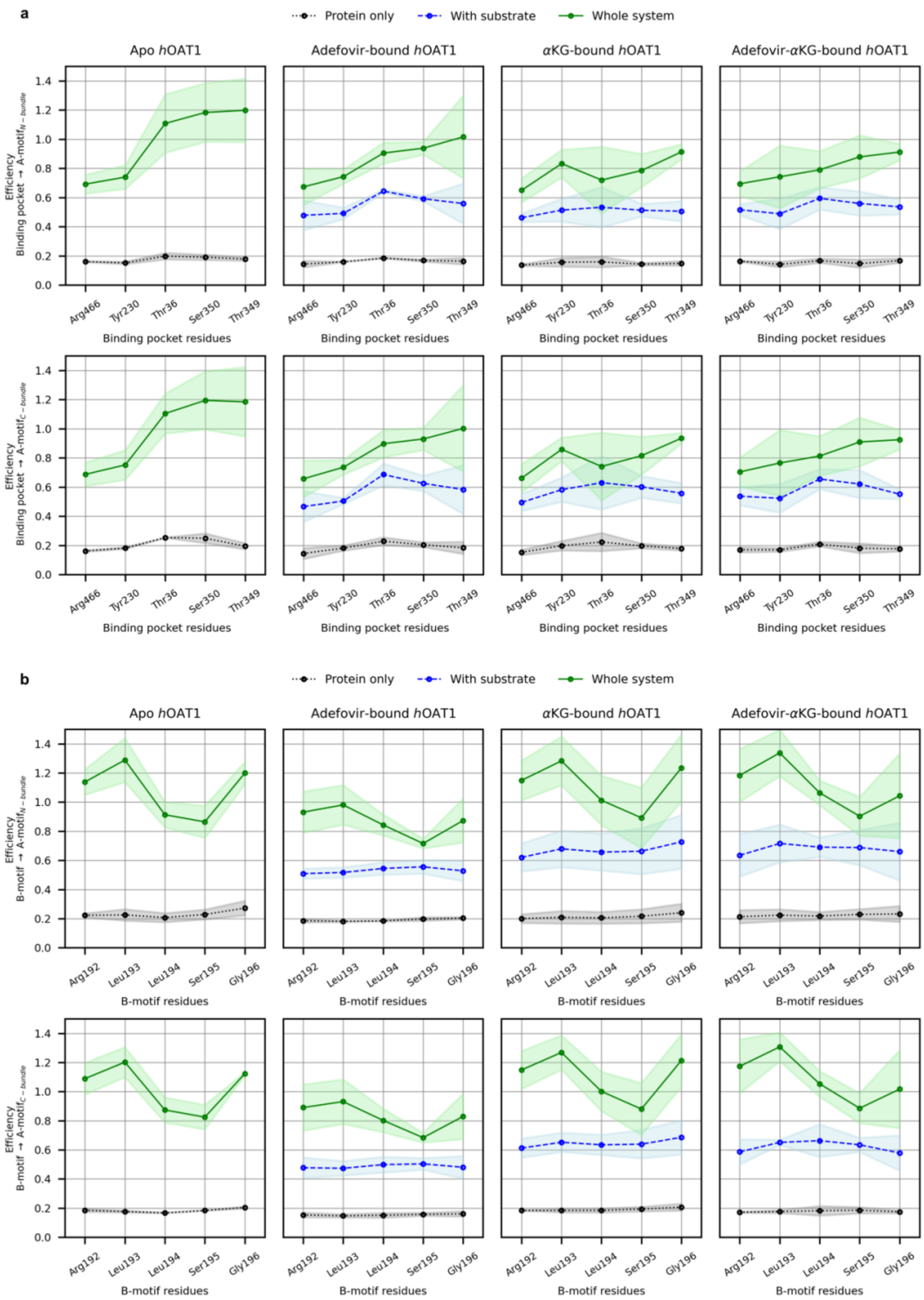


**Figure S7.** Efficiencies of the allosteric pathways from *h*OAT1 (a) inner binding pocket and (b) b-like motif to A-motifs. For each panel, N- and C-bundle efficiencies are provided in top and bottom, respectively. Calculations were performed considering POPC:POPE:Chol (2:1:1) lipid bilayer.

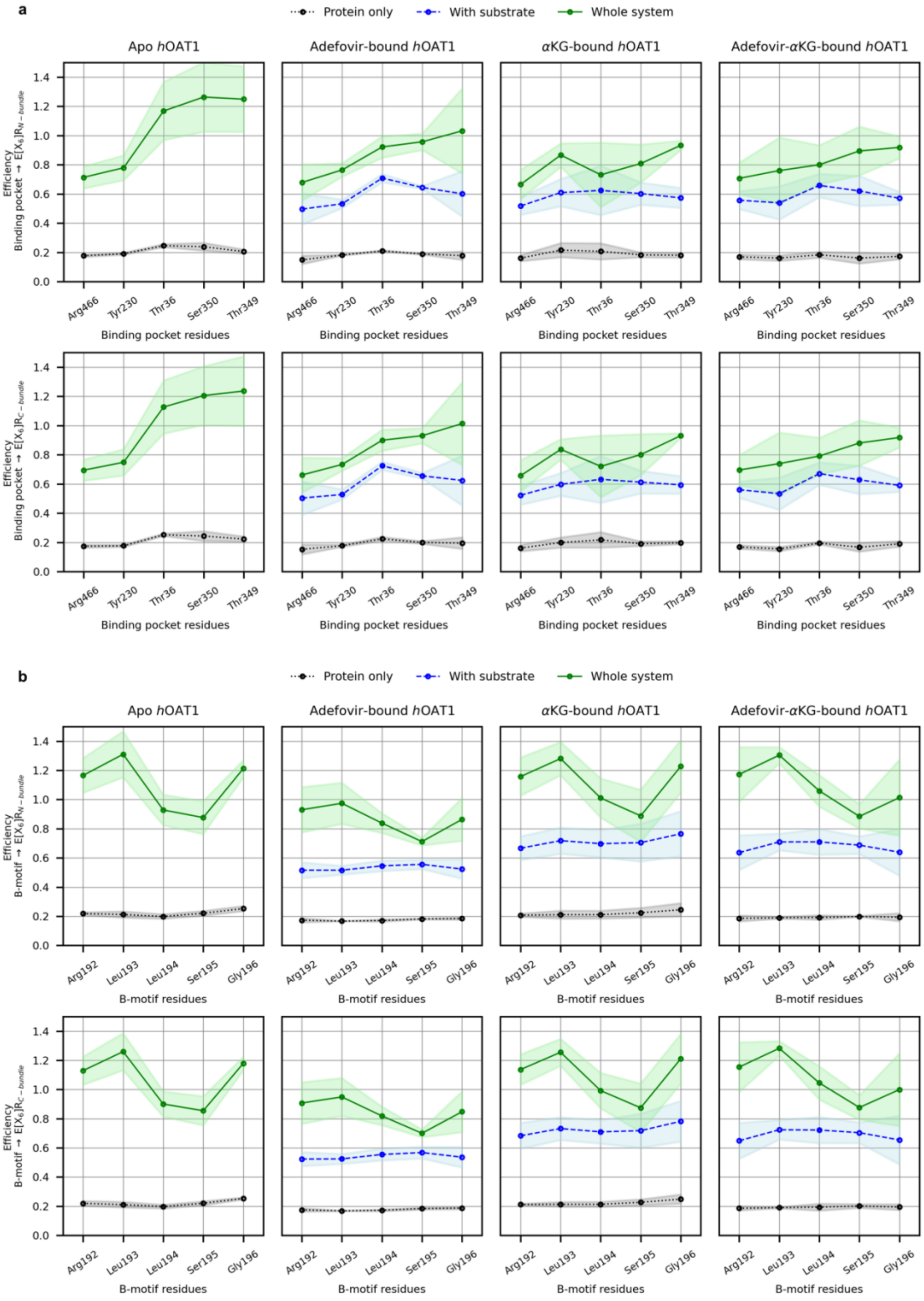


**Figure S8.** Efficiencies of the allosteric pathways from *h*OAT1 (a) inner binding pocket and (b) b-like motif to E[$X_6$]R-motifs. For each panel, N- and C-bundle efficiencies are provided in top and bottom, respectively. Calculations were performed in POPC:POPE:Chol (2:1:1) lipid bilayer.

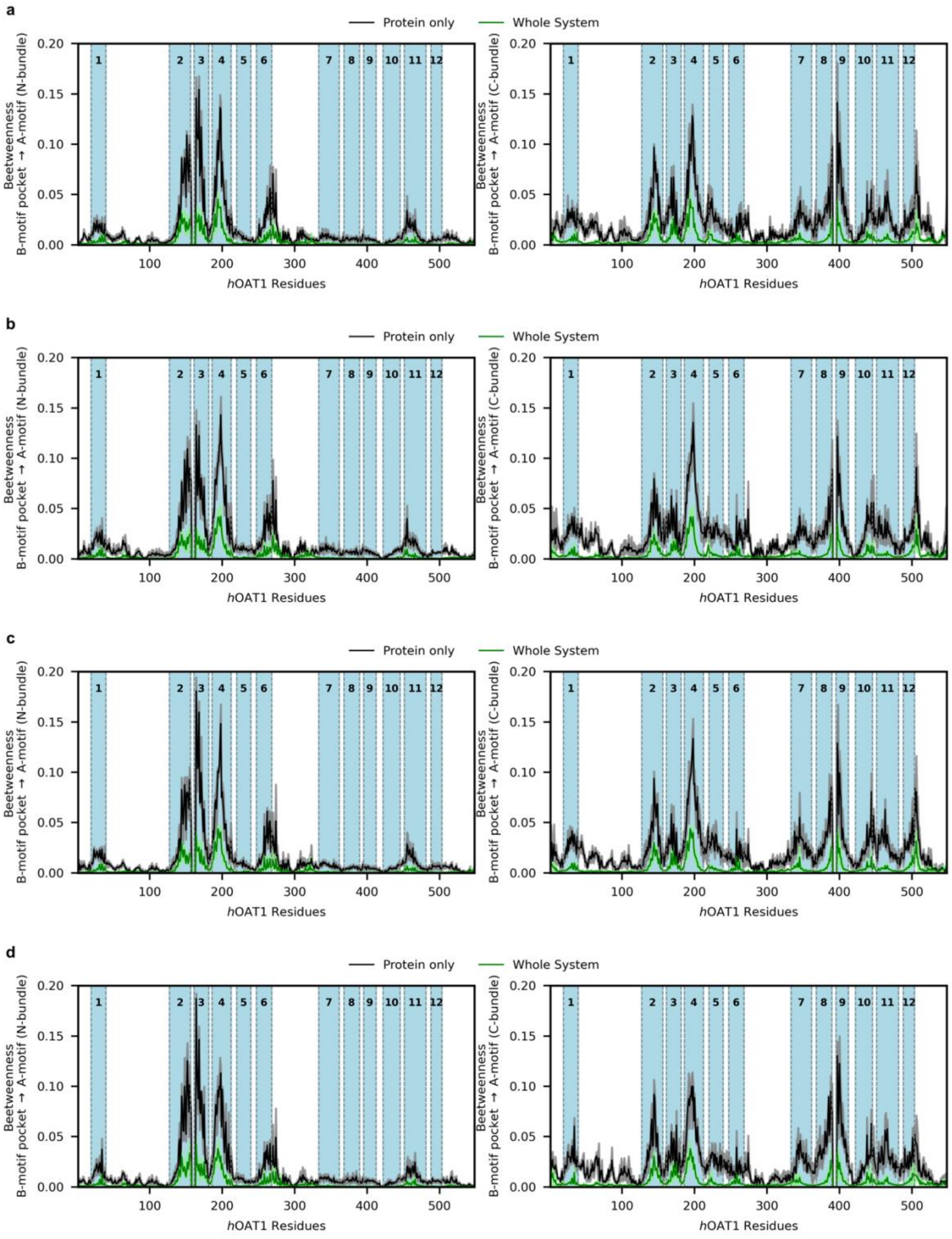


**Figure S9.** Calculated *h*OAT1 per-residue involvement (betweenness) in the allosteric pathway from B-like motif to N-bundle (left) and C-bundle (right) A-motifs for (a) apo, (b) adefovir-bound, (c) $\alpha$KG-bound and (d) adefovir-$\alpha$KG-bound *h*OAT1 systems in POPC:POPE:Chol (2:1:1). TMH topologies are depicted in light blue background.

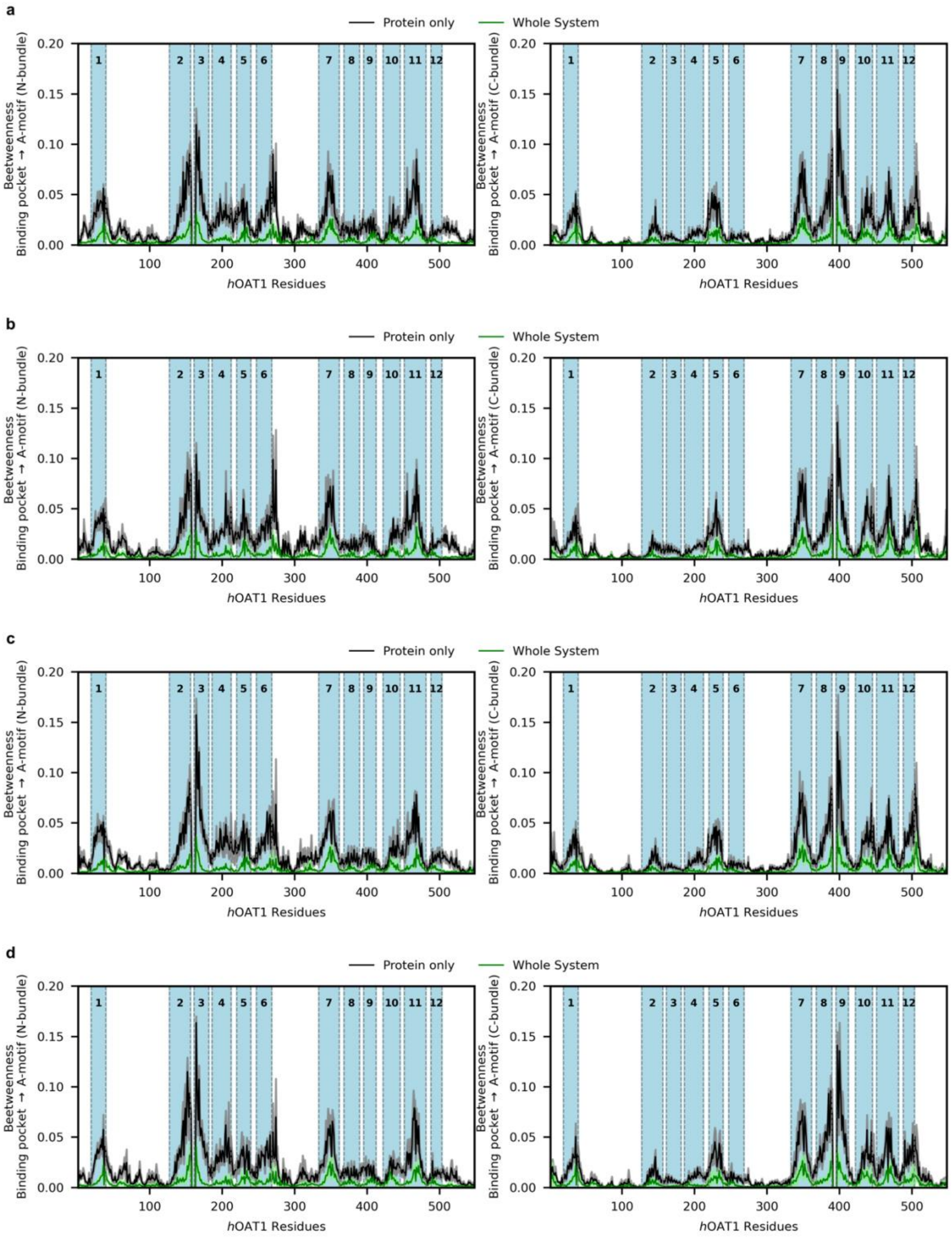


**Figure S10.** Calculated *h*OAT1 per-residue involvement (betweenness) in the allosteric pathway from inner binding pocket to N-bundle (left) and C-bundle (right) A-motifs for (a) apo, (b) adefovir-bound, (c) $\alpha$KG-bound and (d) adefovir-$\alpha$KG-bound *h*OAT1 systems in POPC:POPE:Chol (2:1:1). TMH topologies are depicted in light blue background.

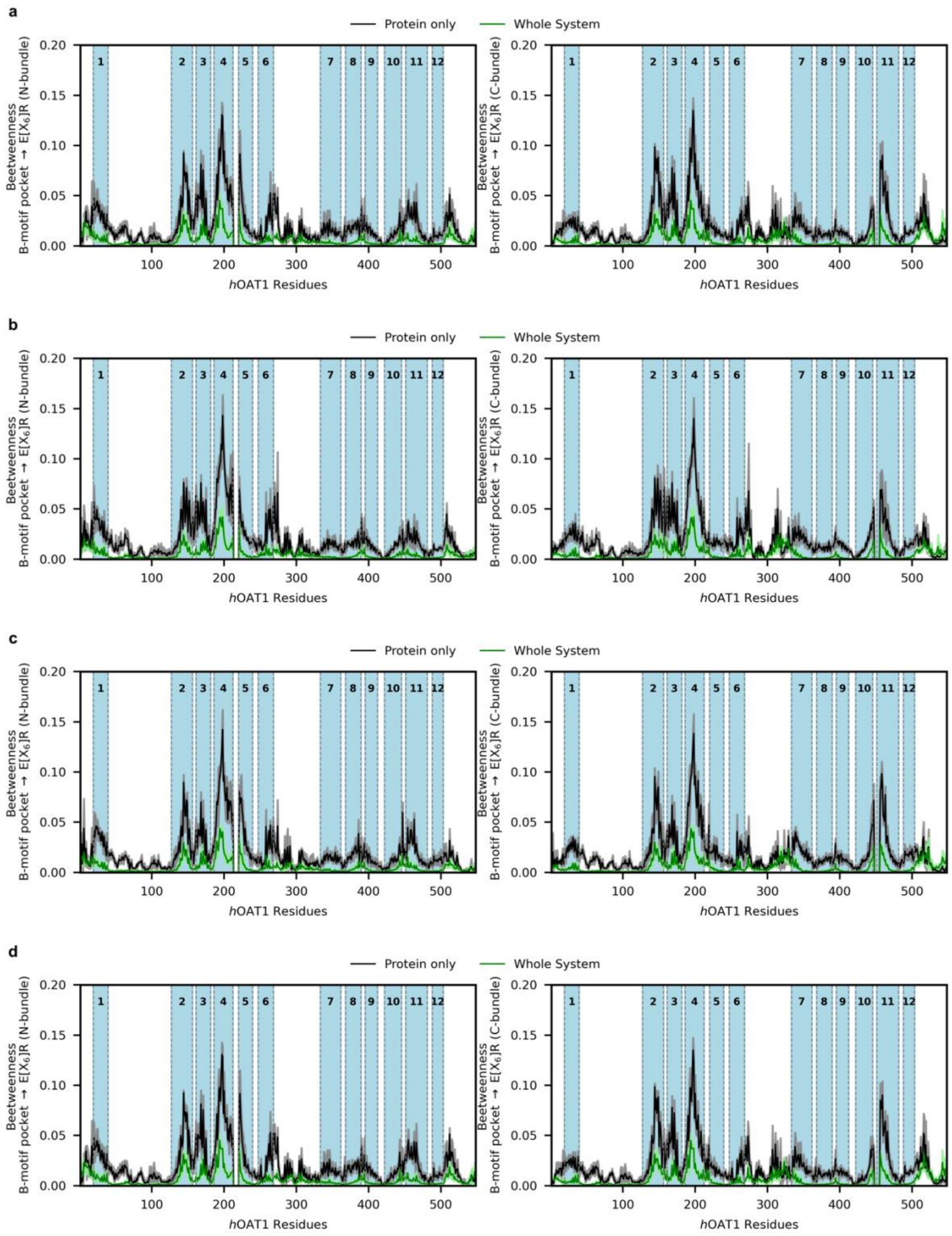


**Figure S11.** Calculated *h*OAT1 per-residue involvement (betweenness) in the allosteric pathway from B-like motif to N-bundle (left) and C-bundle (right) E[$X_6$]R for (a) apo, (b) adefovir-bound, (c) $\alpha$KG-bound and (d) adefovir-$\alpha$KG-bound *h*OAT1 systems. TMH topologies are depicted in light blue background.

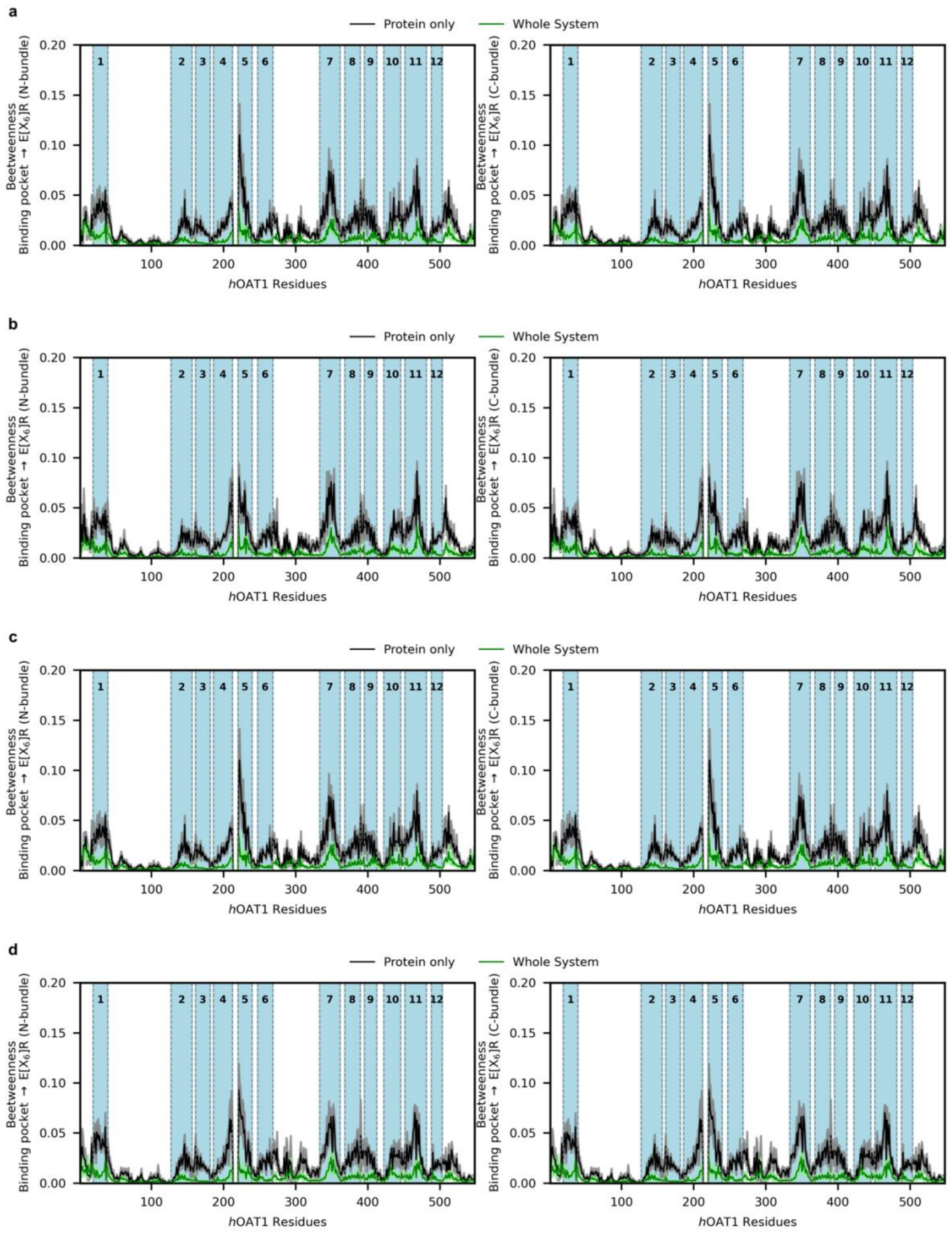


**Figure S12.** Calculated *h*OAT1 per-residue involvement (betweenness) in the allosteric pathway from inner binding cavity to N-bundle (left) and C-bundle (right) E[$X_6$]R for (a) apo, (b) adefovir-bound, (c) $\alpha$KG-bound and (d) adefovir-$\alpha$KG-bound *h*OAT1 systems. TMH topologies are depicted in light blue background.

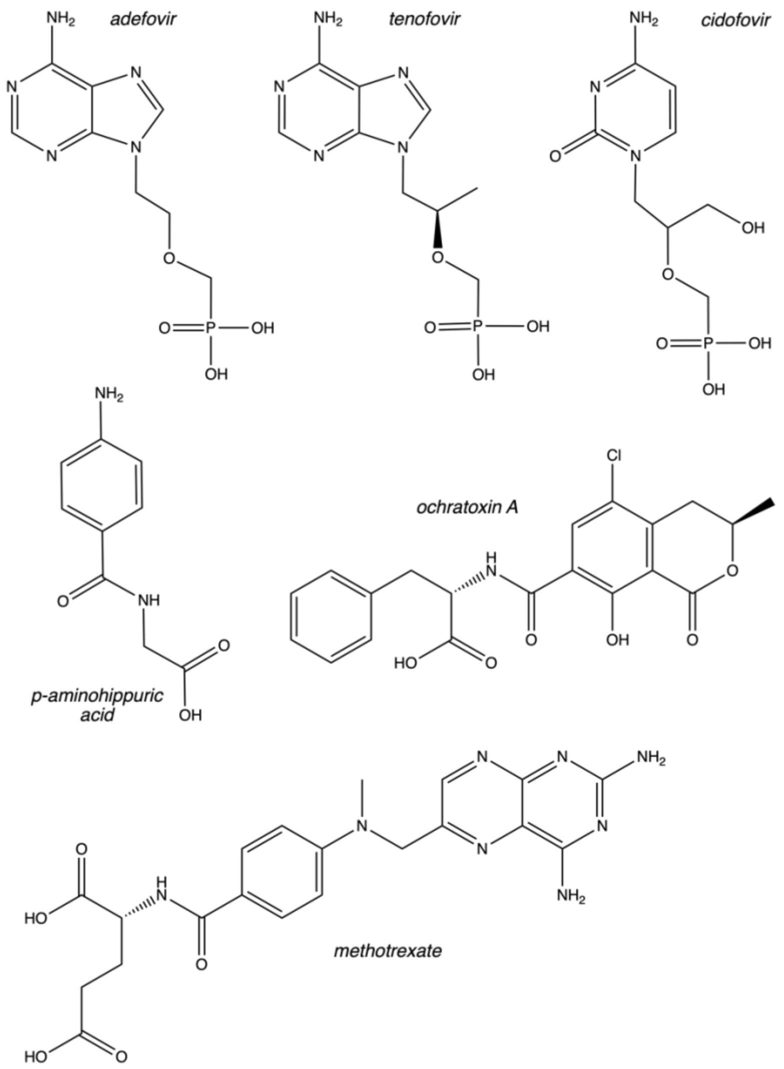


**Figure S13.** Structures of adefovir, tenofovir, cidofovir, ochratoxin A, p-aminohippuric acid, methotrexate.

### 3. Supplemental materials

*Here is place for frcmod and prepi files regarding parametrization of AFV and aKG*

### 4. References


[1] M. Hong, F. Zhou, and G. You, “Critical Amino Acid Residues in Transmembrane Domain 1 of the Human Organic Anion Transporter hOAT1,” *J. Biol. Chem.*, vol. 279, no. 30, pp. 31478–31482, Jul. 2004, doi: 10.1074/jbc.M404686200.

[2] K. Tanaka, W. Xu, F. Zhou, and G. You, “Role of glycosylation in the organic anion transporter OAT1,” *J. Biol. Chem.*, vol. 279, no. 15, pp. 14961–14966, Apr. 2004, doi: 10.1074/jbc.M400197200.

[3] L. Zou *et al.*, “Molecular Mechanisms for Species Differences in Organic Anion Transporter 1, OAT1: Implications for Renal Drug Toxicity,” *Mol. Pharmacol.*, vol. 94, no. 1, pp. 689–699, Jul. 2018, doi: 10.1124/mol.117.111153.

[4] J. L. Perry, N. Dembla-Rajpal, L. A. Hall, and J. B. Pritchard, “A Three-dimensional Model of Human Organic Anion Transporter 1: AROMATIC AMINO ACIDS REQUIRED FOR SUBSTRATE TRANSPORT,” *J. Biol. Chem.*, vol. 281, no. 49, pp. 38071–38079, Dec. 2006, doi: 10.1074/jbc.M608834200.

[5] M. Hong, F. Zhou, K. Lee, and G. You, “The putative transmembrane segment 7 of human organic anion transporter hOAT1 dictates transporter substrate binding and stability,” *J. Pharmacol. Exp. Ther.*, vol. 320, no. 3, pp. 1209–1215, Mar. 2007, doi: 10.1124/jpet.106.117663.

[6] A. N. Rizwan, W. Krick, and G. Burckhardt, “The chloride dependence of the human organic anion transporter 1 (hOAT1) is blunted by mutation of a single amino acid,” *J. Biol. Chem.*, vol. 282, no. 18, pp. 13402–13409, May 2007, doi: 10.1074/jbc.M609849200.

[7] M. Hong, S. Li, F. Zhou, P. E. Thomas, and G. You, “Putative Transmembrane Domain 12 of the Human Organic Anion Transporter hOAT1 Determines Transporter Stability and Maturation Efficiency,” *J. Pharmacol. Exp. Ther.*, vol. 332, no. 2, pp. 650–658, Feb. 2010, doi: 10.1124/jpet.109.160515.

[8] W. Xu, K. Tanaka, A. Sun, and G. You, “Functional Role of the C Terminus of Human Organic Anion Transporter hOAT1,” *J. Biol. Chem.*, vol. 281, no. 42, pp. 31178–31183, Oct. 2006, doi: 10.1074/jbc.M605664200.

[9] A. Janaszkiewicz *et al.*, “Insights into the structure and function of the human organic anion transporter 1 in lipid bilayer membranes,” *Sci. Rep.*, vol. 12, no. 1, p. 7057, Dec. 2022, doi: 10.1038/s41598-022-10755-2.
